# Muscle-specific motor unit firing characteristics in elbow flexors and extensors after cervical spinal cord injury

**DOI:** 10.64898/2026.06.03.729825

**Authors:** Alex T. Benedetto, Sophia T. Jenz, Matthew T. Farley, Bradley S. Heit, James A. Beauchamp, Sina Sangari, Laura McPherson, CJ Heckman, Monica A. Perez, Gregory E.P. Pearcey

**Author notes:** **Correspondence:** Alex T. Benedetto, 3450 N. Lakeshore Drive #2104, Chicago IL, 60657, or, Gregory Pearcey, 230 Elizabeth Avenue, St. John’s, NL, A1C 5S7.

## Abstract

Individuals with cervical spinal cord injury (SCI) often exhibit asymmetric recovery of upper-limb function, with greater weakness in elbow extensors than flexors. To determine whether muscle-specific changes in motor unit (MU) behavior contribute to this disparity, we identified MU firing instants from high-density surface electromyography to characterize MU firing characteristics in the biceps brachii (BIC) and triceps brachii (TRI) of individuals with cervical SCI (n = 20) and non-injured controls (n = 18). We quantified rate-coding behavior and metrics related to persistent inward currents (PICs), including onset-offset hysteresis (ΔF), ascending firing rate nonlinearity, and self-sustained firing. At the group level, BIC MUs in SCI participants showed reduced rate coding and altered ascending firing rate nonlinearity relative to controls. In contrast, TRI MUs showed no clear group-level differences. However, subgroup analysis revealed that SCI participants with low-strength during extension (n = 9) exhibited lower ΔF and longer self-sustained firing durations in TRI MUs than those with high-strength (n = 6). In BIC, SCI participants with low-strength during flexion (n = 8) showed reduced rate-coding behavior relative to high-strength SCI participants (n = 9), with no differences in PIC-related metrics. Together, these results demonstrate muscle-specific alterations in MU firing after cervical SCI that may relate to strength recovery or preservation and underscore the need for nuanced analyses in heterogeneous SCI populations.

**Key points:**

1. Rate coding and nonlinear firing behaviors are significantly altered in the biceps brachii, but not triceps brachii, of participants with cervical spinal cord injury.
2. Strength based subgroup analyses revealed muscle-specific differences in motor unit behaviors that may be associated with strength preservation or recovery following spinal cord injury.
3. Functional heterogeneity following spinal cord injury may mask group differences in motor unit behaviors and warrants careful interpretation of results of future studies.

## INTRODUCTION

Individuals with cervical spinal cord injury (SCI) often exhibit asymmetric recovery of elbow flexors and extensors, with more weakness in the latter (Ditunno *et al.,* 1992; Calancie *et al.,* 2004; McKay *et al.,* 2011; Balbinot *et al.,* 2023). Extensor weakness severely limits functional independence, preventing essential activities such as wheelchair propulsion, transfers, pressure relief, and reaching (Mateo *et al.,* 2015; Javeed *et al.,* 2024). Although an upregulation in reticulospinal projections to the biceps (Sangari & Perez, 2020) and increased intracortical inhibition to the triceps (Butler *et al.,* 2024) may contribute to disparate muscle recovery, further examination of spinal motoneuron dysfunction may provide useful information and illuminate potential therapeutic targets.

Due to the high-fidelity of connections between motoneurons and their innervated muscle fibers, analyses of motor unit (MU) spike times can reveal in-depth information about the firing behavior of spinal motoneurons, which emerges from dynamic combinations of excitatory, inhibitory, and neuromodulatory inputs (Johnson *et al.,* 2017; Chardon *et al.,* 2024). Following SCI, all three of these synaptic inputs can be significantly altered (Holstege C Kuypers, 1987: Hounsgaard *et al*., 1988; Oudega & Perez, 2012), resulting in disordered motor control and involuntary muscle behaviors. Altered MU firing patterns are, therefore, crucial in explaining neuropathophysiological mechanisms underlying muscle-specific imbalances in recovery.

Early studies that examined MU firing behavior after SCI targeted relatively few muscles (Thomas *et al.,* 2014) due to technological limitations. Although recent studies have examined MU behaviors of individuals with SCI in the ankle (Kizyte *et al.,* 2025; Goreau *et al.,* 2026) and wrist musculature (Bao & Lei, 2024; Oliveira *et al.,* 2025), there are limited direct comparisons of biceps brachii and triceps brachii MU firing characteristics in this population (Wiegner *et al.,* 1993). Furthermore, most MU studies in SCI participants have primarily focused on firing rate modulation and variability, recruitment – derecruitment patterns, and twitch force production (for review see Thomas *et al.,* 2014). Although informative, these metrics do not adequately capture and appraise the full interaction of synaptic inputs to motoneurons and the effect of persistent inward currents (PICs), which are the principal mediators of motoneuronal excitability (Lee and Heckman, 1998, 2000).

Recent advances in hardware and software now allow the routine identification of MU spike trains from high-density surface electromyography (HDsEMG), yet there remains a decrement of use in SCI research. Population MU firing patterns are particularly valuable because they allow for the disentanglement of underlying motor commands. Firing rate onset-offset hysteresis (Gorassini and colleagues 1998; 2002) and non-linear firing patterns during the ascending phase (Beauchamp et al., 2023) are complementary quantification techniques that reflect the contribution of PICs to spinal motoneuron output. Combined with realistic simulations of motoneuron firing patterns, these metrics have deepened our understanding of how excitation-inhibition patterns, and neuromodulatory inputs shape motor output (Beauchamp et al., 2023; Chardon et al., 2024), and are potentially valuable tools for understanding pathological MU behavior following SCI.

After SCI, severe damage to the bulbospinal pathways often limits the survival of descending serotonin (5HT) and noradrenaline (NA) fibers below the level of injury, both of which are known to facilitate PICs (Hounsgaard *et al.,* 1988; Heckman and Enoka, 2012). Despite the loss of descending monoaminergic input, PICs are restored to their normal size within months of the injury through the upregulation of constitutively active G-protein coupled receptors, specifically 5-HT_2C_ and NAα_1_ (Murray *et al.,* 2010, 2011; D’Amico *et al.,* 2013; Johnson *et al.,* 2013, Tysseling *et al*., 2017). It is unclear, however, whether the recovery of motoneuronal PICs may contribute to the functional recovery of strength differently in some muscles compared to others.

To identify potential muscle-specific differences in MU firing behavior after SCI, we used identified MU spike instants from HDsEMG to non-invasively quantify MU firing characteristics from the biceps brachii (BIC) and triceps brachii (TRI) of individuals with a cervical level SCI. We first explored group-level differences between SCI and non-injured (NI) control participants for each muscle. We then sub-grouped the SCI individuals based on muscular strength to further investigate muscle-specific behaviors that may be linked to imbalanced functional recovery.

## METHODS

### Participants

Twenty individuals with chronic SCI (≥1 year post injury; mean injury duration 10.5 ± 25.6; mean age 37.6 ± 10.1 years; 3 females) and eighteen neurologically intact (NI) control individuals (mean age 34.1 ± 9.48 years; 2 females) participated in this study. All participants provided written informed consent, and study procedures were approved by the Northwestern University Institutional Review Board in accordance with the Declaration of Helsinki, except registration. For individuals in the SCI group, neurological level of injury was determined using the International Standards for Neurological Classification of Spinal Cord Injury (ISNSCI) examination, with all injuries at or above C8. SCI participants were classified by the American Spinal Cord Injury Impairment Scale (AIS) as AIS A (n=3), AIS B (n=2), AIS C (n = 9) and AIS D (n = 6). All participants reported no history of musculoskeletal injury in the right upper limb at the time of testing and were not taking any monoaminergic altering medications. Participants in the SCI group were not instructed to abstain from the use of prescribed anti-spasmodic or pain medications (e.g., Baclofen or Gabapentin) prior to testing, but their use was noted for post hoc investigation of a possible correlation with MU behaviors, and none were found. A full list of participant demographics related to the study can be found in Table 1.

**Table 1.** SCI Participant Demographics. Abbreviations: AIS = American SCI Impairment Scale (Grade A, B, C, or D); Gab = Gabapentin; Bac = Baclofen; T = traumatic; NT = non-traumatic.

| Participant | Age | Sex | Time post injury (yrs) | AIS | Level of Injury | Medication | Etiology |
| --- | --- | --- | --- | --- | --- | --- | --- |
| SCI01 | 33 | M | 9.25 | C | C8 | Bac | T |
| SCI02 | 49 | F | 16.1 | B | C5 | None | T |
| SCI03 | 38 | M | 1.22 | C | C5 | Gab | T |
| SCI04 | 43 | M | 3.28 | D | C5 | None | T |
| SCI05 | 30 | M | 2.25 | D | C4 | Bac/Gab | T |
| SCI06 | 26 | M | 3.61 | A | C6 | None | T |
| SCI07 | 26 | M | 7.5 | C | C4 | None | NT |
| SCI08 | 22 | F | 4.33 | C | C4 | Bac/Gab | NT |
| SCI09 | 45 | F | 117 | D | C6 | Bac/Gab | NT |
| SCI10 | 40 | M | 12.7 | D | C7 | None | T |

| Participant | Age | Sex | Time post injury<br>(yrs) | AIS | Level of Injury | Medication | Etiology |
| --- | --- | --- | --- | --- | --- | --- | --- |
| SCI11 | 50 | M | 1.81 | C | C4 | None | T |
| SCI12 | 30 | M | 0.97 | C | C7 | Gab | T |
| SCI13 | 32 | M | 1.11 | A | C4 | Bac | T |
| SCI14 | 48 | M | 1.8 | C | C7 | Bac/Gab | T |
| SCI15 | 57 | M | 1.39 | D | C4 | Bac/Gab | T |
| SCI16 | 49 | M | 1.57 | A | C6 | None | T |
| SCI17 | 27 | M | 2.31 | B | C6 | Bac/Gab | T |
| SCI18 | 29 | M | 1.47 | C | C4 | Bac | T |
| SCI19 | 46 | M | 2.71 | D | C5 | None | T |
| SCI20 | 33 | M | 11.5 | C | C7 | None | T |

### In session protocol

All participants were seated in a custom-built experiment chair placed in front of a monitor to provide visual feedback of their elbow torque production as shown in Fig 1A. None of the participants had notable asymmetries, so the right arm was tested in everyone and was secured to a custom device with an attached calibrated ATO load cell (model ATO-LCC-DYMH-103). The limb was positioned at 90° shoulder flexion and 90° elbow flexion with forearm fully supinated and straps placed at the wrist and elbow to minimize extraneous movement. Additional trunk support was provided with an over-the-shoulder chest strap. Each session began with participants performing three trials of 3-second maximal voluntary isometric elbow flexion and extension contractions (MVCs). One minute of rest was given between each trial, and verbal encouragement was provided to ensure maximal effort. If maximum torque continued to increase, additional trials were collected until three stable values were obtained. Maximal voluntary contraction torque (TQ_MVC_) was quantified by averaging the maximum value from these three trials and used to normalize the subsequent experimental, submaximal contractions. For these contractions, participants performed triangular flexion and extension (10 second ascending phase; 10 second descending phase) to a peak amplitude of 30% their TQ_MVC_. Participants were instructed to do their best to stay within the target boundary, which had a range of ±2% TQ_MVC_. A minimum of 2 trials were performed in each direction, but trials were discarded and re-done if the torque profile did not adequately match the visual target. Target contraction shapes for MVC and submaximal ramps are depicted in Fig 1A.

**Figure 1.**
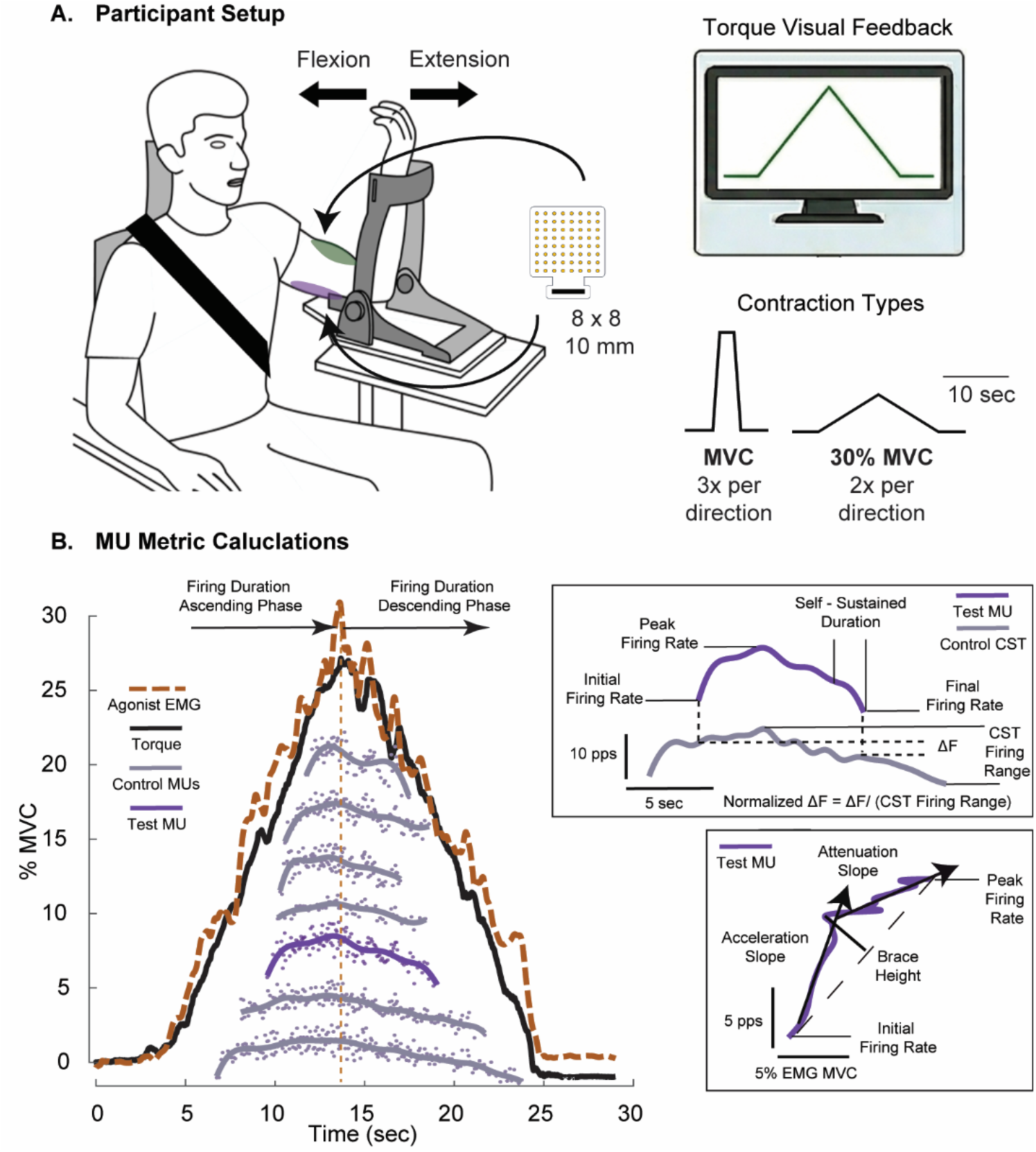
Experimental methods. (A) Participant setup in custom-built experimental chair and isometric elbow torque force transducer. Illustration of high-density surface EMG electrode, electrode placement locations at the biceps brachii (BIC; green) and triceps brachii (TRI; purple), and the isometric contraction shapes for MVC and 30% MVC. (B) Data processing workflow from smoothed instantaneous discharge rates to calculated MU metrics. Insert boxes show the main MU metric calculations using the purple MU as an example. For the ΔF calculation, the gray CST is calculated using the firing rates of all the gray MUs in the ramp trial. The purple MU is also used for the Brace Height example.

### High-density surface electromyography

High-density surface electromyography (HDsEMG) electrode grids (64 electrodes, 8 × 8 configuration, 10 mm inter-electrode distance, GR10MM0808, OT Bioelettronica, Turin, Italy) were placed over the midline of the muscle belly of the biceps brachii (BIC) and triceps brachii (TRI). HDsEMG signals were recorded in monopolar mode, sampled at 2048 Hz, amplified 150 times, and band-pass filtered between 10–500 Hz using a signal amplifier (OT Bioelettronica, Turin, Italy).

### Data Analysis

#### Task Performance

The quality of the triangular contraction was assessed visually and by quantifying torque accuracy, steadiness, and cross correlation to the provided visual target. Torque was sampled at 2048 Hz and filtered offline using a lowpass 50Hz fifth-order Butterworth filter. Torque accuracy was quantified using the root mean squared error (RMSE) of deviations from the visual target provided. Torque samples that remained within the boundary were given an error score of 0, while deviations outside the boundary received an error score equal to the distance of the deviation on a normalized %MVC scale. Error scores were then used to calculate the RMSE for the trial. Torque steadiness during segments of the ascending and descending phases of the contraction were calculated using coefficient of variation (CoV) of the detrended torque signal. Due to the ±2% TQ_MVC_ boundary window and slight timing variations in participants initiating the ascent and descent phases, we assessed the middle 8 seconds of each phase rather than the full 10 seconds. This allowed for consistent analysis across participants without penalizing for behavioral timing of each phase. The Pearson coefficient for the Torque-to-Target cross correlation of the entire contraction was quantified using the *xcorr* function in MATLAB to account for lag.

#### Peak EMG and Muscle Coactivation

The magnitude of EMG was quantified in both the agonist and the antagonist muscle for each submaximal ramp contraction and MVC trial. After applying a single differential to the monopolar signals across the 64-channel grid, each single-differential signal was band-pass filtered between 20-500 Hz, rectified, and then low-pass filtered at 4 Hz for envelope visualization. Signal-to-noise ratios were used to identify the 30 channels from the grid with the best signal quality. These signals were then averaged to obtain one global signal per muscle for each trial and MVC, which was used for further analysis. This method allowed for more consistent analysis across participants with varying numbers of noisy channels during the experimental session. EMG during maximal voluntary contraction (EMG_MVC_) was calculated from the MVC trials using a 1-second sliding average window to identify the maximum EMG amplitude for each MVC contraction, and these values were used to normalize the EMG envelopes obtained during the submaximal triangular contractions. Peak EMG of the agonist muscle during the triangular contractions was also calculated using a 1-second sliding window. Muscle coactivation of the agonist and antagonist muscles was quantified during the same 1-second window (i.e., at peak agonist activation) using a coactivation index: Antagonist / (Antagonist + Agonist) (Thomas *et al.,* 1998). This was calculated using the EMG_MVC_ normalized envelopes and is reported on a scale from 0 -1, where higher values indicate a greater amount of antagonist coactivation. Additionally, we assessed the overall shape of the agonist EMG envelope using the *xcorr* MATLAB function to obtain the Pearson coefficient for an EMG-to-Target cross correlation.

#### EMG decomposition

EMG signals were visually inspected and removed if substantial artifacts or noise were present. The remaining signals were decomposed into individual MU spike trains using a blind source separation algorithm, and manual editing was performed on the decomposed spike trains to correct for any minor errors (Negro et al., 2016, Del Vecchio et al., 2020). Custom MATLAB scripts were used to calculate the instantaneous firing rates as the inverse of the interspike intervals, and the spike trains were smoothed for further analysis using support vector regression using hyperparameters described previously (Beauchamp *et al*., 2022).

#### Analysis of motor unit firing characteristics

Although typically reported as a percentage of TQ_MVC_, recruitment and derecruitment thresholds were quantified as a percentage of EMG_MVC_ to better account for the magnitude of muscle activation and the variety of torque and EMG envelope relationships across participants. Initial, peak and final firing rates were calculated and used to quantify firing rate modulation ranges. Self-sustained duration quantifies the percentage of MU firing duration that extends beyond the expected firing duration. This value is calculated by subtracting the ascending firing duration from the descending firing duration and is normalized to the total firing duration (Afsharipour *et al*., 2020; Hassan *et al.,* 2021). The split between the ascending and descending phase is for each trial was determined using the peak of agonist EMG envelope during the triangular contraction (refer to fig 1B).

The contribution of persistent inward currents (PICs) to prolonged firing and firing rate acceleration was estimated by quantifying firing rate hysteresis and non-linearity, two hallmark PIC characteristics that result in prolonged output from, and amplification of, synaptic input. Firing rate hysteresis is typically calculated using a paired MU analysis technique (i.e., Delta frequency, or ΔF) which has been validated and widely used over the last two decades (Gorassini *et al*., 2002; Mesquita *et al.,* 2024). This method quantifies the onset-offset hysteresis of a higher threshold “test” MU as the difference in firing rate of a lower threshold “reporter” MU at the instants of recruitment and derecruitment of the test MU. Our modified approach replaced individual reporter MUs with a cumulative spike train (CST) constructed from all MUs except the test MU (as shown in Fig 1B) in an attempt to provide a better estimate of synaptic input to the MU pool. We applied the same inclusion criteria for the calculation that is applied to the unit-wise approach: (1) The test unit must be recruited at least 1s after the reporter unit or CST to ensure full PIC activation (Bennett *et al*., 2001; Powers *et al*., 2008); (2) the test–reporter pair received common synaptic input, indicated by a rate–rate correlation of ≥ 0.7 (Gorassini *et al*., 2004); and (3) the reporter unit exhibited a firing range of at least 0.5 pulses per second (pps) while the test unit was active (Hassan *et al*., 2021). To further control for SCI and NI group differences in MU firing rate modulation, ΔF was normalized to the maximum possible value by dividing the ΔF by the difference in CST modulation from test MU recruitment to the lowest firing rate.

Firing rate non-linearity was quantified using a geometric approach to identify the maximum orthogonal deviation of the ascending MU firing rate, known as Brace Height (Beauchamp *et al*., 2023). To calculate Brace Height in this study, the smoothed MU firing trace was plotted as a function of the agonist EMG envelope during the contraction (Gomes, *et al.,* 2024) and the deviation from a theoretical linear increase from firing rate at recruitment to peak firing rate was determined. To account for scaling due to differences in rate modulation, Brace Height values are normalized to the height of a right triangle (%rTri), where the hypotenuse represents recruitment to peak MU firing. The Brace Height calculation also identifies two key phases of the ascending firing rate modulation, the Acceleration and Attenuation slopes (pps/%MVC), which can provide further insight into patterns of excitation-inhibition coupling and neuromodulatory drive (Figure 1B). All MUs were manually inspected for irregularities and removed from further analysis if their brace height was greater than 100 %rTri, acceleration phase exceeded 2 seconds in duration or had a negative slope, or their firing rate peaked after the peak of the EMG envelope (Beauchamp *et al*., 2023).

#### Motor unit tracking

We identified repeated MUs across trials to reduce potential bias in the statistical analysis due to repeated observations. This was done by estimating the MU action potential (MUAP) waveforms with spike-triggered averaging and computing a 2-D cross-correlation between the spatial representation of the MUAPs between trials (Martinez-Valdes *et al.,* 2017; Del Vecchio *et al.,* 2019). Mus were determined to be repeated (i.e., tracked) if their correlation coefficient exceeded 0.75, and all uniquely identified MUs were assigned distinct identifiers (UnID).

### Statistical Analysis

We used linear mixed-effects models to examine the effects of group, muscle and their interaction on each task performance variable and MU property, following the general structure Outcome ∼ Group*Muscle + Covariates + (1+Muscle|Participant). Categorical predictors were coded using sum contrasts (±0.5; e.g., Muscle: BIC = 0.5, TRI = −0.5; Group: SCI = 0.5, NI = −0.5) so that model intercepts represented grand means and fixed effects reflected average differences across levels. Continuous covariates were mean-centered to reduce non-essential multicollinearity and improve interpretability of interaction terms and were retained in the final model when they significantly improved fit (assessed with the *anova* function. The model was fit using the *lme4* package (Bates *et al*., 2015) in R (R Core Team, 2021) and assumptions were evaluated using the *Performance* package (Lüdecke *et al*., 2021). The Group × Muscle interaction was evaluated by comparing nested models with and without the interaction term using likelihood ratio tests with the *anova* function. The interaction was retained in the final model when it significantly improved model fit. Estimated marginal means and Group × Muscle contrasts were calculated using the *emmeans* package in R (Lenth, 2023). General outcome-specific data transformations and model modifications are detailed below.

#### Task performance metrics

In the task performance models, Muscle was replaced with Direction (Flexion vs. Extension), and we included age and sex as covariates when significant as both factors are known to influence maximal force output and steadiness (Tracy *et al*., 2005; Inglis & Gabriel, 2021; Yamaguchi *et al*., 2023). Trial number was also included as a covariate to account for potential improvements with task repetition. An additional fixed effect of Phase (ascending vs. descending) was included in the torque CoV model to investigate potential Group × Phase, Direction × Phase, or Group × Direction × Phase interactions.

Residual diagnostics from our initial models indicated deviations from normality and variance for several of the variables. EMG-to-Target and Torque-to-Target Pearson cross correlation coefficients were transformed using Fisher’s r-to-z transformation (*atanh*) prior to modeling. Given that correlation coefficients are bounded and possess non-linear variance properties, this transformation yields an approximately normal sampling distribution with stabilized variance, consistent with assumptions of linear mixed-effects modeling. Model assumptions were re-evaluated following transformation through visual inspection of residual and Q–Q plots, confirming improved normality and homoscedasticity. Estimated marginal means were back-transformed to the correlation scale for reporting, and standard errors and confidence intervals on the r scale were calculated using the delta method. Statistical tests and contrasts were performed on the Fisher z scale to account for non-linearity of the transform and preserve standardized model effect size comparisons.

#### Group-level comparisons of MU firing characteristics

All models for the MU metrics initially included MU recruitment threshold, age, and sex as covariates based on known associations with MU firing properties (Hassan *et al., 2021,* Beauchamp *et al.,* 2023, Jenz *et al.,* 2023). Due to the observed variation in EMG patterns across participants, the inclusion of peak EMG and EMG-to-Target correlation (i.e., EMG envelope shape) as covariates was also explored. If any of these covariates did not significantly improve model fit, they were removed from the final model. In addition to the participant random intercept with muscle slope, a random intercept for the unique MU identifier (1 | UnID) was also included to avoid treating repeated observations of the same unit as independent. For the acceleration slope and attenuation slope metrics, a log transform was applied to improve residual diagnostics. Estimated marginal means were back transformed to their original scale for visualization and interpretability, with the group contrasts reported as the ratio of geometric means.

We further investigated the number of MUs with higher ascending rate modulation compared to descending rate modulation using a binomial generalized linear mixed-effects model with a logit link. The model estimated the probability that a MU had higher ascending than descending rate modulation, with MUs coded as 1 when ascending modulation exceeded descending modulation and 0 otherwise. Fixed effects included group and muscle, with sex and EMG-to-Target cross correlation entered as additional predictors. Random intercepts for participant and UnID accounted for clustering of MUs within individuals.

#### Relationship between maximum voluntary torque output and MU firing characteristics

Due to the substantial strength variability in our SCI participants, we further examined the relationship between strength and MU/task performance metrics by grouping the individuals based on their TQ_MVC_ values. Cluster analysis was performed using k-means clustering. Observations were partitioned into two clusters (k = 2) based on Euclidean distance, with cluster membership determined by minimizing within-cluster variance around the cluster centroids. Cluster validity was evaluated using silhouette analysis, yielding an average silhouette value of 0.58. Cluster 1 contained all NI male values for both muscles, while the 2 NI females were placed in cluster 2 for both muscles. The SCI males were split between clusters 1 and 2, and all 3 SCI females were placed in cluster 2. Due to the low sample size of NI females and their cluster separation from the NI males, we only included male participants in the final analysis. Although this is slightly limiting, it is not unreasonable given ∼78% of individuals who suffer from traumatic SCI are males (National Spinal Cord Injury Statistical Center, 2024). We performed a second cluster analysis with only males to ensure proper grouping, and the clusters remained the same with a silhouette value of 0.59. For the BIC, 9 of the SCI males were grouped into cluster 1 with all 16 NI males, and the remaining 8 SCI males were grouped into cluster 2. For the TRI, 6 SCI males were placed in cluster 1 with all 16 NI males, and the remaining 7 SCI males were grouped in cluster 2. From here on, the three cluster-based subgroups are referred to as NI, high-strength SCI, and low-strength SCI. It should be noted that three of the SCI participants were classified as high-strength BIC (cluster 1) but low-strength TRI (cluster 2).

The MU metric differences in strength groups for each muscle were explored separately using linear mixed effects models with this general formula: lmer(Metric ∼ Subgroup + covariates + (1|Participant) + (1|UnID)). As previously described, covariates were removed if they did not improve model fit. For the task performance metrics, the model was simplified to lmer(Metric ∼ Subgroup + covariates + (1|Participant)). For MVCs, linear fixed effects models were used to estimate differences between the group-cluster subgroups for each contraction direction/muscle: lm(MVC ∼ Subgroup + Age).

## RESULTS

### Age of participants

There was no significant difference in age between SCI and NI groups for BIC data (NI: 34.1 ± 9.5 years, n = 18; SCI: 37.6 ± 10.1 years, n = 20; Welch’s t(35.94) = −1.13, p = 0.27) or TRI data (NI: 34.1 ± 9.5 years, n = 18; SCI: 39.0 ± 9.8 years, n = 15; Welch’s t(29.3) = −1.45, p = 0.16).

### Torque output during MVC

The linear-mixed effects model revealed a significant Group × Direction interaction (X^2^ = 7.74, p = 0.005), with the SCI group producing significantly lower torque than the NI group for both flexion (28.6 ± 5.09 Nm vs. 51.2 ± 5.46 Nm, p < 0.001, d = 1.95) and extension (21.1 ± 5.49 Nm vs. 60.4 ± 5.46 Nm, p < 0.001, d = 3.40), as shown in figure 2A. Despite lower estimated marginal means during extension, there was no significant difference between directions within the SCI group (p=0.08, d = 0.65), but the NI group had significantly higher extension values compared to flexion (p = 0.02, d = 0.8). On average, males produced higher torque than females (p < 0.001), while age (p = 0.85) was unrelated.

**Figure 2.**
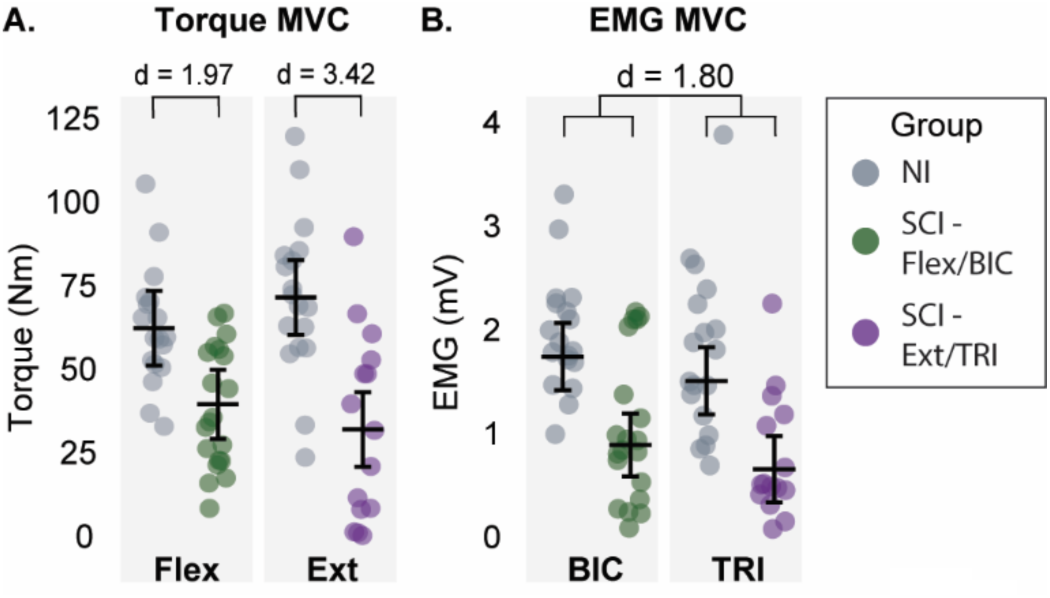
Torque and EMG during Maximal voluntary contractions. (A) Participant torque values obtained during maximal voluntary elbow ffexion (Flex) and extension (Ext). Each data point is a single participant value with error bars indicating estimated group means and S5% CI from the linear model analysis. Brackets and effect size (Cohen’s d) indicate significant group contrast. (B) Similar plot but for maximal agonist EMG recorded during maximal voluntary contractions. BIC values were recorded during isometric elbow ffexion (Flex), and TRI values were recorded during isometric elbow extension (Ext). SCI Flex/BIC: n = 20, SCI Ext/TRI: n = 15, NI Flex/BIC: n = 18 NI Ext/TRI: n = 18.

### Muscle activity during MVC

EMG (mV) measured during the MVC trials revealed significantly lower muscle activation in the SCI group compared to the NI group (0.823 ± 0.15 vs. 1.674 ± 0.15; p < 0.001, d = 1.80). The estimated marginal means are as follows: SCI BIC: 0.939 ± 0.15 NI BIC: 1.79 ± 0.16; SCI TRI: 0.706 ± 0.16 NI TRI: 1.56 ± 0.16. There was an overall muscle effect, with BIC having significantly higher values than TRI (p = 0.05, d = 0.49), but there was no significant Group × Muscle interaction (p = 0.6). Age was significant (p = 0.006) with younger individuals having higher values, but sex was not significant (p = 0.054, males: 1.49 ± 0.09 mV, females: 1.01 ± 0.23 mV).

#### Task Performance Metrics

##### Error in torque

Torque accuracy, quantifying deviation from the visual target using a root mean squared error (RMSE), showed a significant Group × Direction interaction (X^2^ = 8.34, p = 0.004), and pairwise contrasts revealed significantly greater error during extension for the SCI group compared to the NI group (0.89 ± 0.12 % MVC vs. 0.43 ± 0.12 % MVC; p = 0.008, d = 1.97). However, flexion error was similar between the groups (SCI: 0.63 ± 0.06 % MVC, NI: 0.56 ± 0.06 % MVC; p = 0.42).

##### Variability in torque

The linear mixed effects model for the coefficient of variability (CoV) in torque did not reveal any significant interactions between Group, Direction, or Phase (Ascending or Descending). However, there was a significant Phase effect with greater CoV in the Descending phase across all participants and directions (p < 0.001). The SCI group showed increased CoV compared to the NI group, but it did not reach statistical significance (p = 0.06). The group estimated marginal means for each combination of Direction and Phase are as follows: Ascending Extension (SCI: 4.98 ± 0.65%, NI: 4.18 ± 0.66%), Ascending Flexion (SCI: 3.99 ± 0.29%, NI: 3.20 ± 0.31%), Descending Extension (SCI: 5.96 ± 0.65%, NI: 5.16 ± 0.66%), and Descending Flexion (SCI: 4.98 ± 0.29% NI: 4.18 ± 0.31%).

##### Torque-to-Target correlation

The linear mixed effects model revealed a significant Group × Direction interaction (X^2^ = 9.62, p = 0.002). Pairwise contrasts showed that the SCI group had significantly lower cross-correlation values than the NI group for extension (0.992 ± 0.001 vs. 0.995 ± 0.0004 (Fisher back-transformed); p = 0.001, d = 1.76 z-scale), but not flexion (0.994 ± 0.001 vs. 0.994 ± 0.001 (Fisher back-transformed); p = 0.7 z-scale).

##### Peak EMG during triangular contractions

There was a significant Group × Direction interaction (X^2^ = 5.64, p = 0.02) for peak agonist EMG, with pairwise contrasts revealing significantly higher amplitudes in the SCI group compared to the NI group for TRI during extension (26.1 ± 1.82 % MVC vs. 17.1 ± 1.67 % MVC; p = 0.001, d = 2.98) but not for BIC during flexion (18.9 ± 1.95 %MVC vs. 19.0 ± 2.05 %MVC; p = 0.95).

##### Muscle Coactivation

There was a significant Group × Direction interaction (X^2^ = 6.81, p = 0.009) for agonist – antagonist muscle coactivation. Pairwise contrasts found significantly higher antagonist muscle activity during flexion in the SCI group compared to the NI group (0.49 ± 0.05 vs. 0.25 ± 0.05; p = 0.001, d = 6.5), but there was no significant difference in coactivation between the groups during extension (SCI: 0.15 ± 0.03 vs. NI: 0.16 ± 0.03; p = 0.88).

##### EMG-to-Target correlation

The linear mixed effects model revealed a significant Group × Direction interaction (X^2^ = 6.46, p = 0.01). Pairwise contrasts found the SCI group to have significantly lower cross-correlation values than the NI group for TRI during extension (0.973 ± 0.006 vs. 0.990 ± 0.002 (Fisher back-transformed); p = 0.002, d = 2.84 z-scale), but values for BIC during flexion were similar (0.970 ± 0.005 vs. 0.974 ± 0.004 (Fisher back-transformed); p = 0.58 z-scale).

#### Motor Unit Metrics

##### Group-level comparisons of motor unit firing characteristics

###### Motor unit identification

For the TRI, we were unable to identify MUs from one SCI participant and four others were excluded due to their inability to reasonably perform the extension task. Altogether, 1,209 MU spike trains were analyzed in this study (SCI BIC: n = 338, SCI TRI: n = 262, NI BIC: n = 301, NI TRI: n = 308). The number of identified MUs varied across participants, but the average number identified per trial was similar across groups for both muscles (SCI BIC: 8.45 ± 4.57, SCI TRI: 8.73 ± 5.56, NI BIC: 8.36 ± 4.87, NI TRI: 8.56 ± 6.44). Of the 1,209 units, 866 were identified as unique MUs, with 343 (∼40%) MUs identified in both trials for each muscle within a given participant. Again, the average number of tracked units per trial was similar across groups for both muscles (SCI BIC: 4.15 ± 3.15, SCI TRI: 5.86 ± 3.57, NI BIC: 4.94 ± 4.04, NI TRI: 5.50 ± 6.02). The average number of decomposed MUs per trial for each participant is listed in table 2.

**Table 2.**
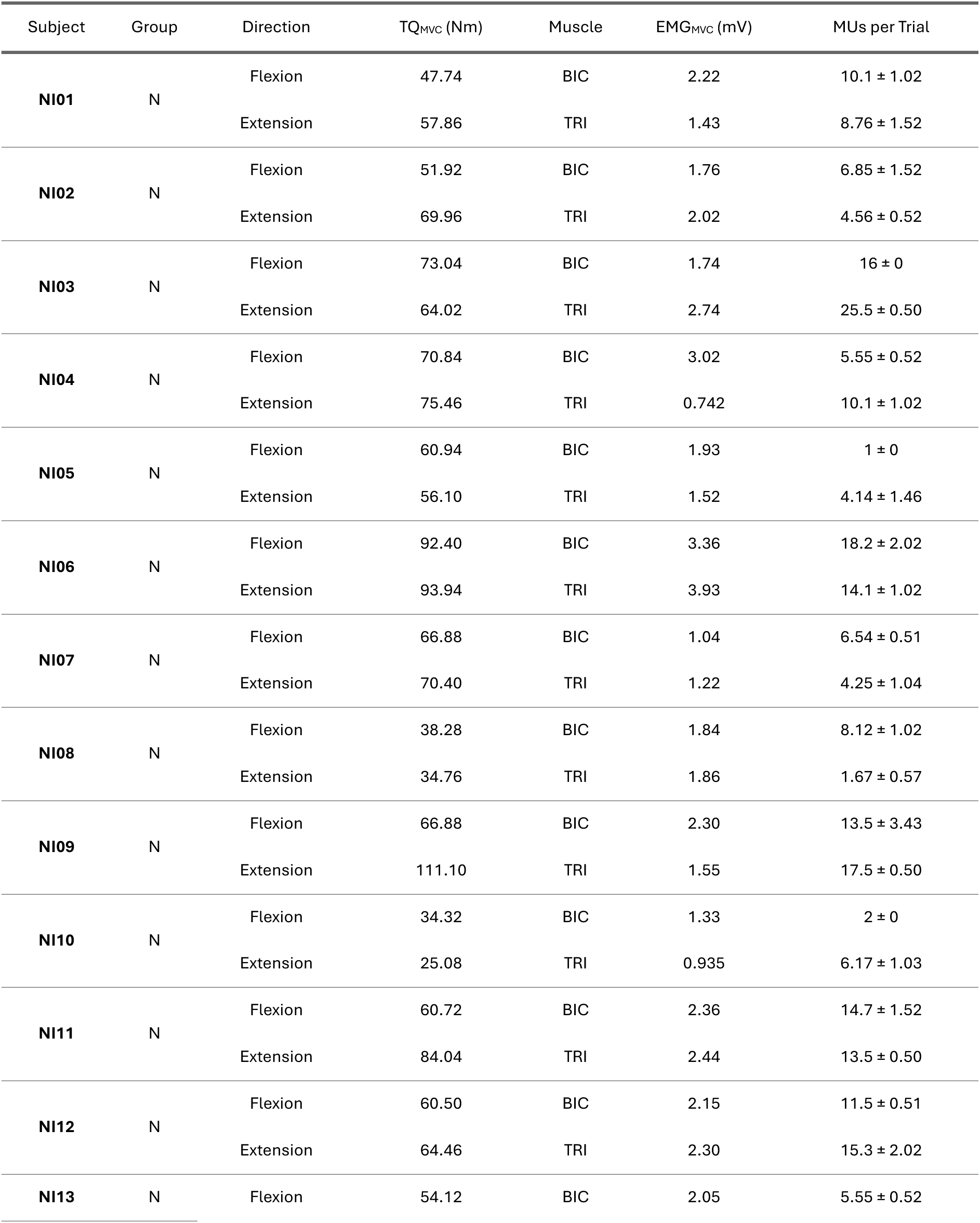
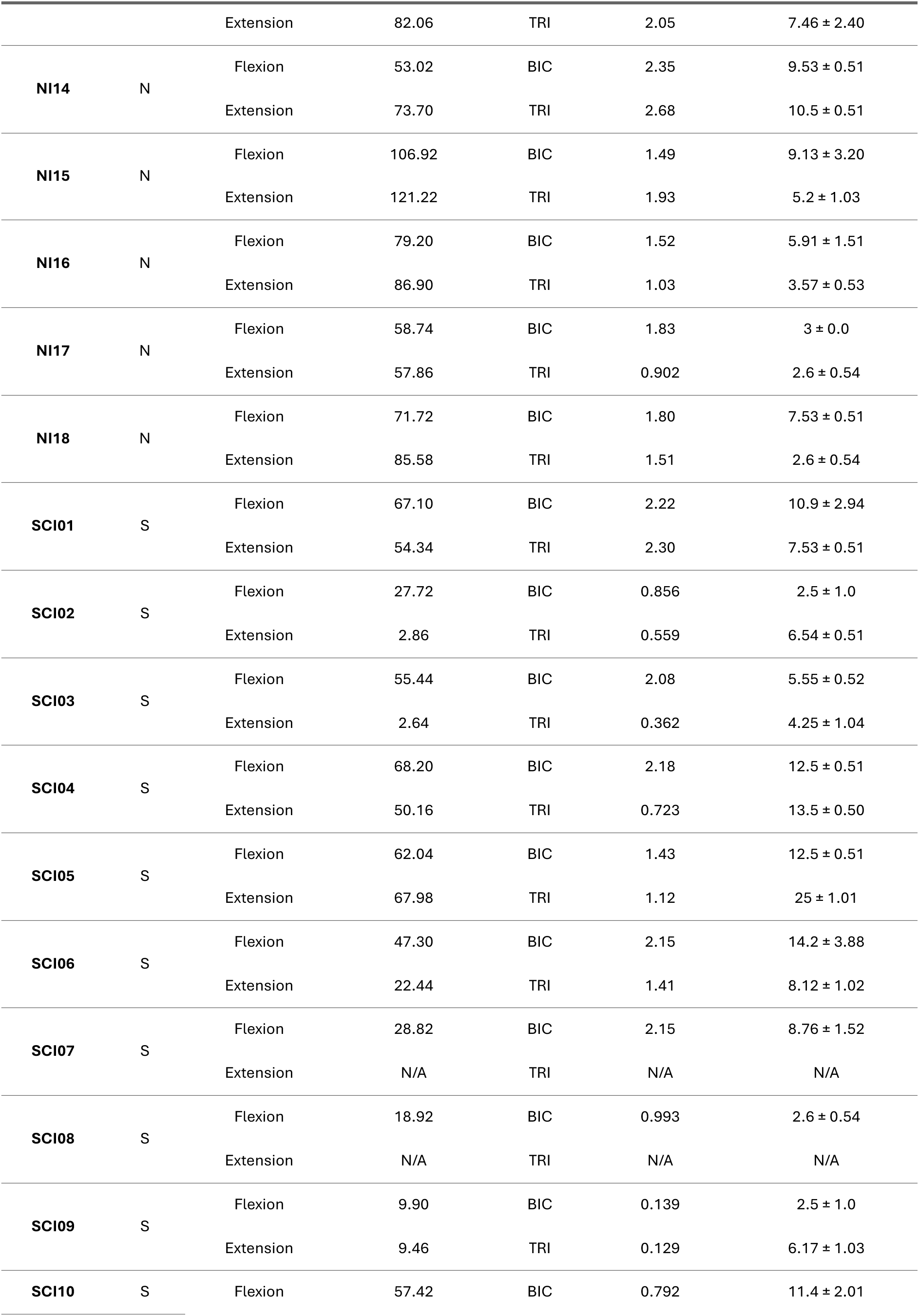
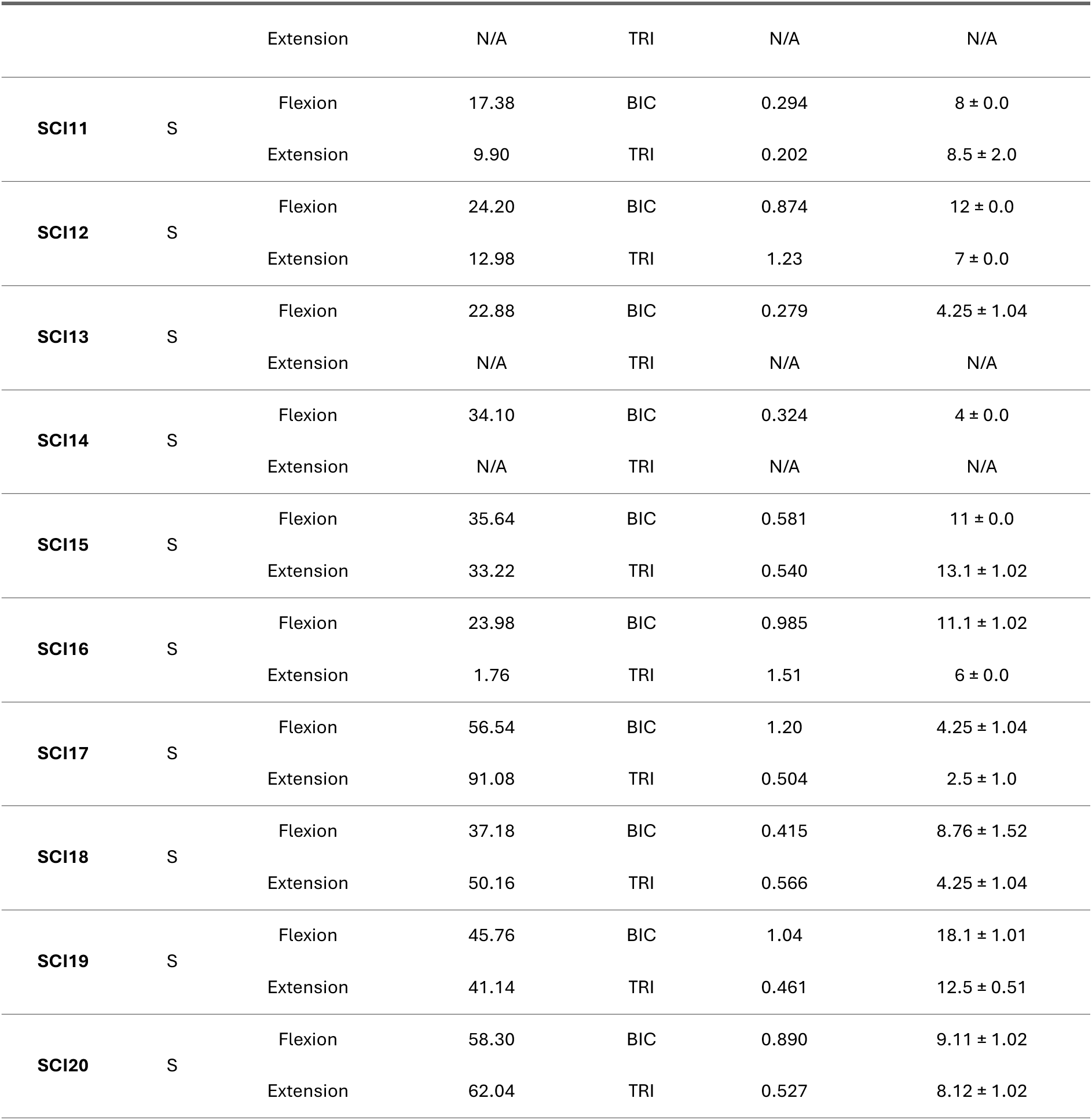
Maximal voluntary contraction and motor unit yield for each participant.

The linear-mixed effects model revealed there was no significant Group × Muscle interaction (p = 0.06), effect of muscle (p = 0.9), or effect of group (p = 0.08). The estimated means for each group are as follows: SCI BIC: 12.1 ± 1.15 %EMG_MVC_, NI BIC: 9.88 ± 1.18 %EMG_MVC_; SCI TRI: 12.3 ± 1.02 %EMG_MVC_, NI TRI: 10.1 ± 0.96 %EMG_MVC_. The recruitment threshold distribution and binned counts are shown in figure 3.

**Figure 3.**
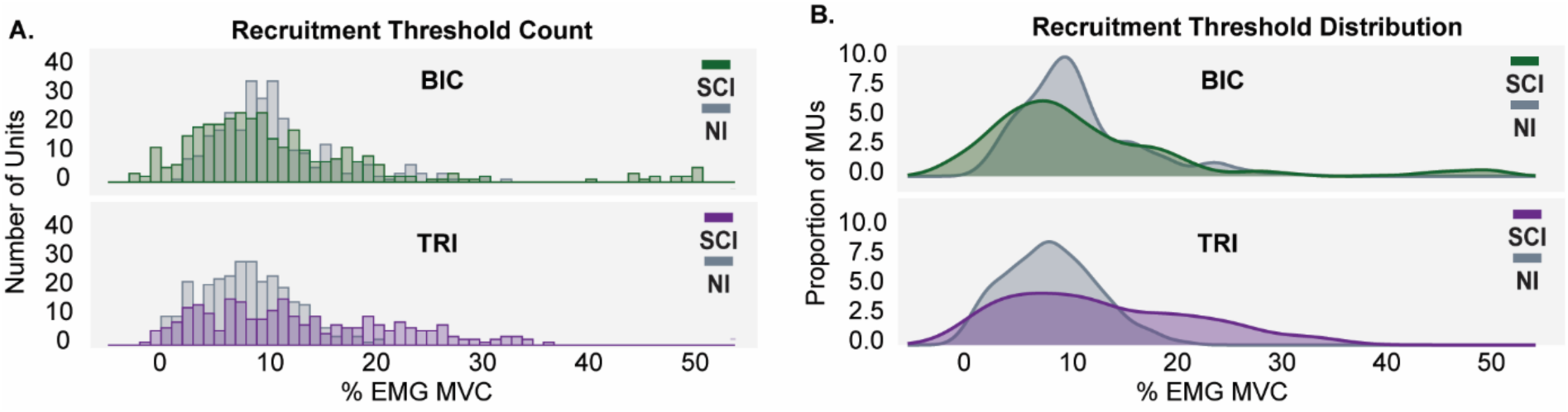
Recruitment threshold count and distribution. (A) recruitment thresholds calculated as % EMG_MVC_ binned in 1% increments to show total number of MUs at that contraction level. (B) recruitment thresholds plotted as a continuous distribution of total proportion of MUs at a given contraction level. For each plot, SCI data are colored and NI data are in gray. SCI BIC: n = 20; SCI TRI: n = 15; NI BIC: n = 18; NI TRI: n = 18.

###### Initial Firing Rate

The linear-mixed effects model revealed a significant group effect (p = 0.003, d = 0.71) with rates (pps) significantly reduced in the SCI group compared to the NI group across both muscles (SCI = 7.96 ± 0.40; NI = 9.8 ± 0.41). There was also a significant muscle effect (p < 0.001), but there was no significant Group × Muscle interaction (p = 0.4). The estimated marginal means for each group are as follows: SCI BIC: 8.57 ± 0.42; NI BIC: 10.4 ± 0.44; SCI TRI: 7.34 ± 0.42; NI TRI: 9.19 ± 0.42.

###### Peak Firing Rate

There was a significant Group × Muscle interaction for peak firing rate (X^2^ = 11.03 p < 0.001). The SCI group had significantly lower firing rates in the BIC compared to the NI group (14.2 ± 0.58 pps vs. 17.8 ± 0.60 pps, p < 0.001, d = 1.82), but there was no difference between the groups in the TRI (SCI: 14.2 ± 0.74 pps vs. NI: 14.6 ± 0.73 pps, p = 0.67).

###### Final Firing Rate

There was also a significant Group × Muscle interaction for final firing rate (X^2^ = 5.65, p = 0.017). Again, the SCI group had significantly lower firing rates in the BIC compared to the NI group (7.15 ± 0.35 vs. 8.20 ± 0.36, p = 0.04, d = 0.43), but there was no group difference in the TRI (SCI: 7.04 ± 0.30 vs. NI: 6.8 ± 0.29, p = 0.56).

###### Ascending Firing Rate Modulation

There was a significant Group × Muscle interaction (X^2^ = 11.02, p < 0.001) for ascending rate modulation. The SCI group had significantly lower ascending rate modulation compared to the NI group in the BIC (6.70 ± 0.31 pps vs. 8.51 ± 0.33 pps; p < 0.001, d = 0.66) but not the TRI (7.63 ± 0.58 pps vs. 6.66 ± 0.55 pps; p = 0.24).

###### Descending Firing Rate Modulation

Although there was no significant Group × Muscle interaction (p = 0.54) or muscle effect (p = 0.13), there was a significant group effect (p = 0.001, d = 0.6) with the SCI group having lower values on average across muscles (7.03 ± 0.47 pps vs. 8.80 ± 0.48 pps).

###### Total Firing Rate Modulation

The linear mixed model revealed a significant Group × Muscle interaction (X^2^ = 6.39, p = 0.011), with the SCI group having significantly lower rate modulation than the NI group for the BIC (8.18 ± 0.48 pps vs. 10.47 ±0.50 pps; p < 0.001, d = 0.92). There were no significant group differences for the TRI (SCI: 8.82 ± 0.60 pps vs. NI: 8.84 ± 0.59 pps, p = 0.97).

###### Ascending Dominant Rate Modulation

There was no significant Group × Muscle interaction (p = 0.9) or effect of muscle (p = 0.14). However, there was a significant main effect of group (SCI/NI contrast odds ratio = 1.66 ± 0.31; p = 0.006, d = 0.28), indicating a higher probability of SCI MUs (0.46 ± 0.05) exhibiting greater ascending than descending rate modulation compared to NI MUs (0.34 ± 0.04).

**Figure 4.**
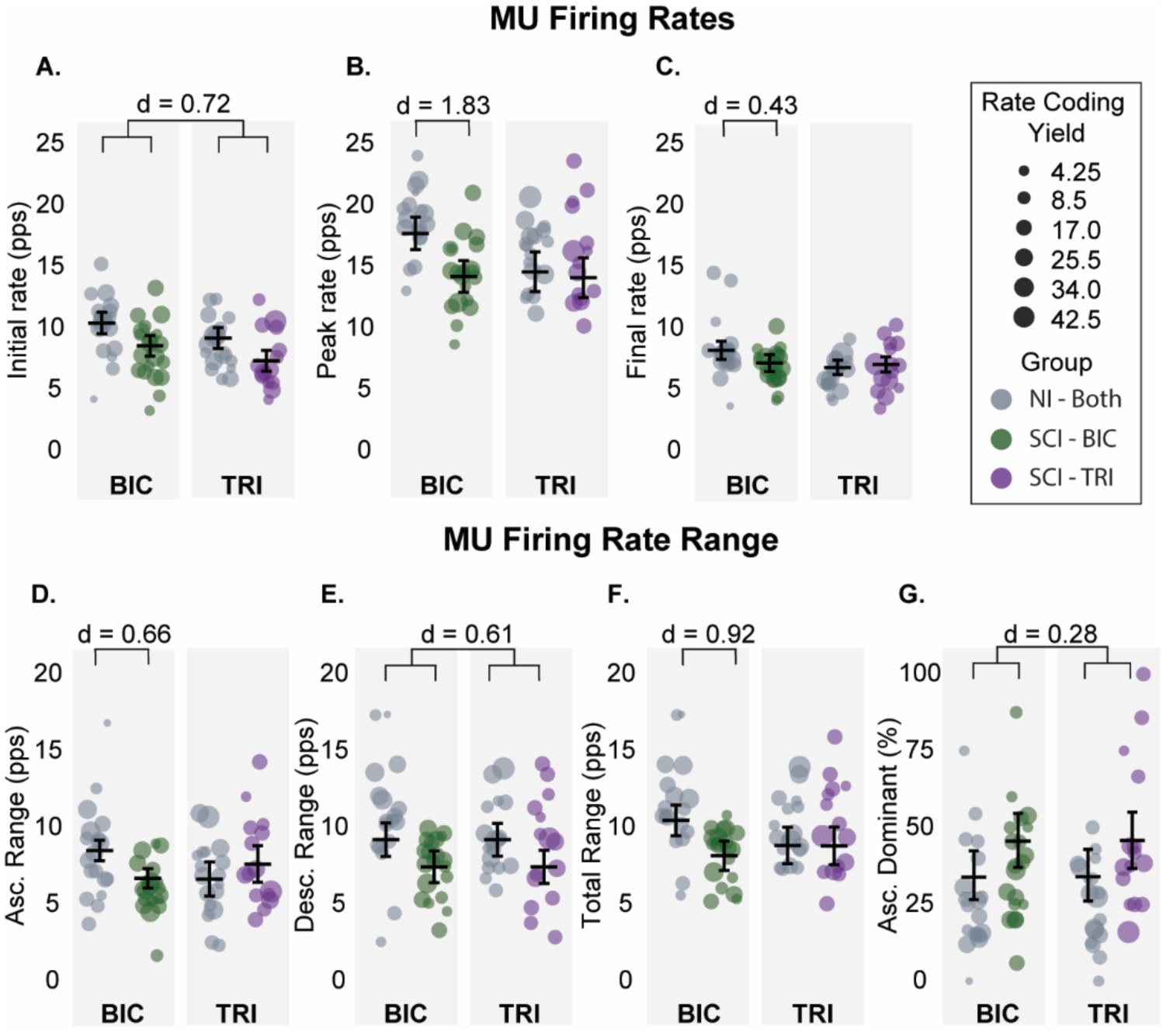
Motor unit (MU) Rate-Coding Metrics. (A-C) Instantaneous MU firing rates calculated at time of recruitment (initial rate) (A), maximum firing rate (peak rate) (B), and derecruitment (final rate) (C). (D-F) MU firing rate modulation ranges calculated from values A-C. (D) Ascending rate modulation (Asc. Range) is the range from initial (A) to peak (B) firing rate. (E) Descending rate modulation (Desc. Range) is the range from peak (B) to final (C) firing rate. (F) Total rate modulation is the minimum firing rate to maximum firing rate. (G) Ascend Dominant is the probability that a MU will have greater ascending rate modulation than descending rate modulation. For A-G, each data point represents the median-aggregated MU data for each participant, with size reflecting the yield for each observation (larger data points contribute more data to the model), and brackets with effect size (Cohen’s d) indicate significant contrasts. Plots A-F show the estimated marginal means and 95% CI from the LMER results; Plot G shows the estimated probability and S5% CI from a binomial GLMER. SCI BIC: n = 20; SCI TRI: n = 15; NI BIC: n = 18; NI TRI: n = 18.

###### Onset-offset hysteresis

There was no significant Group × Muscle interaction for ΔF (p = 0.8), and neither group (p = 0.6) nor muscle (p = 0.29) were significant predictors. The estimated marginal means for each group are as follows: SCI BIC: 4.31 ± 0.40 pps; SCI TRI: 3.79 ± 0.43 pps; NI BIC: 4.56 ± 0.42 pps; and NI TRI: 4.04 ± 0.40 pps. When examining normalized ΔF, there was also no significant Group × Muscle interaction (p = 0.6), group (p = 0.30) or muscle (p = 0.55) effect. The estimated marginal means for each group are as follows: SCI BIC: 0.383 ± 0.03; NI BIC: 0.345 ± 0.03; SCI TRI: 0.408 ± 0.03; NI TRI: 0.370 ± 0.03.

###### Self-sustained firing duration

There was no significant Group × Muscle interaction (p = 0.3) or muscle effect (p = 0.57) but the SCI group showed significantly greater (p = 0.043, d = 0.42) self-sustained firing duration than the NI group on average across muscles (20.6 ± 3.2% vs. 11.1 ± 3.2%). The estimated marginal means for each group are as follows: SCI BIC: 21.7 ± 3.8%; NI BIC: 12.2 ± 4.1%; SCI TRI: 19.4. ± 4.0%; NI TRI: 9.92 ± 3.8%.

**Figure 5.**
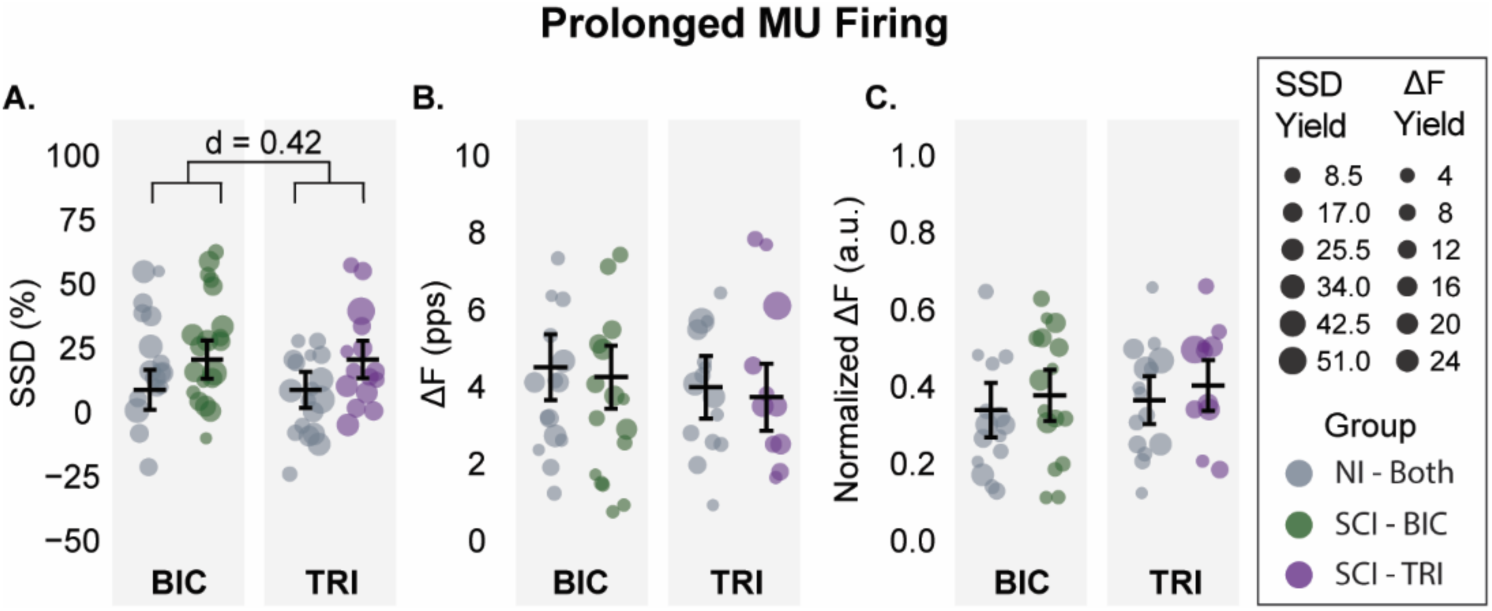
Prolonged motor unit firing metrics. (A) Self-sustained firing duration (SSD) calculated as the percentage of MU firing duration that exceeds the expected total firing duration. (B) Firing rate hysteresis (ΔF or ΔFrequency) calculated using a modified paired-unit analysis technique with a cumulative spike train (CST) in place of the “control unit”, and (C) firing rate hysteresis normalized to the firing rate range of the CST while the “test unit” is active. Each data point represents the median-aggregated MU data for each participant. Data point size reflects the yield for each observation (larger data points contribute more data to the model). Sizes vary because SSD and ΔF were not calculated for every MU due to exclusion criteria. Means and 95% CI estimated from the LMER results. SCI BIC: n = 20; SCI TRI: n = 15; NI BIC: n = 18; NI TRI: n = 18.

###### Brace height

The linear mixed effects model for brace height revealed a significant Group × Muscle interaction (X^2^ = 4.06, p = 0.044). The SCI group had significantly higher values than the NI group in BIC (0.443 ± 0.02 %rTri vs. 0.358 ± 0.02 %rTri; p = 0.003, d = 0.65), but TRI values were similar between groups (SCI: 0.515 ± 0.02 %rTri vs. NI: 0.499 ± 0.01 %rTri; p = 0.47).

###### Acceleration duration

The linear mixed effects model for acceleration phase duration did not have a significant Group × Muscle interaction (p = 0.5), but there was significant effect of muscle (p < 0.001) and of group (p = 0.01, d = 0.44). Durations (sec) were longer in the SCI group compared to the NI group (1.48 ± 0.05 vs. 1.28 ± 0.05), and the estimated marginal means are as follows: SCI BIC: 1.34 ± 0.06; NI BIC 1.13 ± 0.06; SCI TRI: 1.63 ± 0.07; NI TRI 1.42 ± 0.06.

###### Acceleration slope

The linear mixed effects model for acceleration slope did not have a significant Group × Muscle interaction (p = 0.6), group effect (p = 0.21), or muscle effect (p = 0.61). The estimated marginal means for the slopes (pps/%MVC) are as follows: SCI BIC: 2.12 ± 0.30; NI BIC 2.34 ± 0.35; SCI TRI: 2.01 ± 0.33; NI TRI 2.22 ± 0.35).

###### Attenuation slope

There was a significant Group × Muscle interaction for attenuation slope (X^2^ = 11.75, p < 0.001). In the BIC, slopes (pps/%MVC) were significantly reduced in the SCI group compared to the NI group (0.28 ± 0.10 vs. 0.44 ± 0.11; p = 0.02, d = 0.58). However, slopes in the TRI were not significantly different (p = 0.5) between the SCI group (0.29 ± 0.11) and the NI group (0.25 ± 0.10).

**Figure 6.**
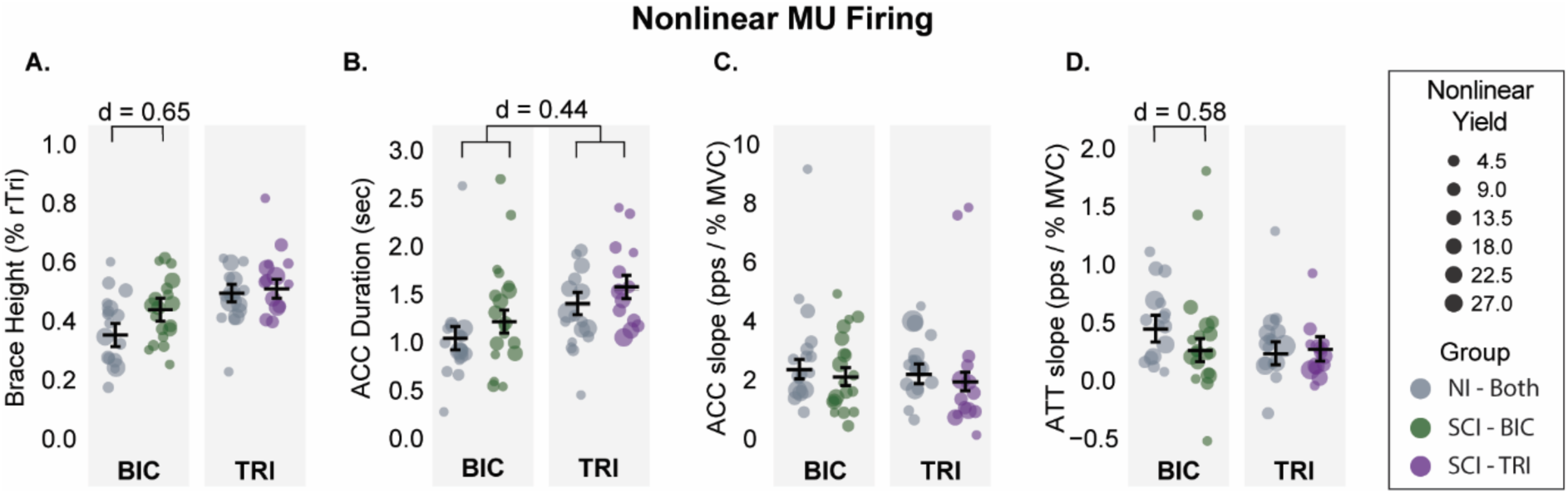
Nonlinear motor unit firing metrics. (A) Brace Height calculated as the maximum orthogonal deviation from a theoretical linear firing rate from initial to peak firing rate. (B) Duration of the firing rate acceleration (ACC) phase (secondary firing rate) from initial firing rate to firing rate saturation at the attenuation phase (tertiary firing rate). (C) Firing rate slope during the acceleration (ACC) phase. (D) Firing rate slope during the attenuation (ATT) phase. Each data point represents the median-aggregated MU data for each participant. Data point size reflects the yield for each observation (larger data points contribute more data to the model). Means and 95% CI estimated from the LMER results (C and D estimated means are back transformed from the log-scale to their original scale for visualization). Brackets and effect size (Cohen’s d) indicate significant group contrast. SCI BIC: n = 20, SCI TRI: n = 15, NI BIC: n = 18, NI TRI: n = 18.

**Figure 7.**
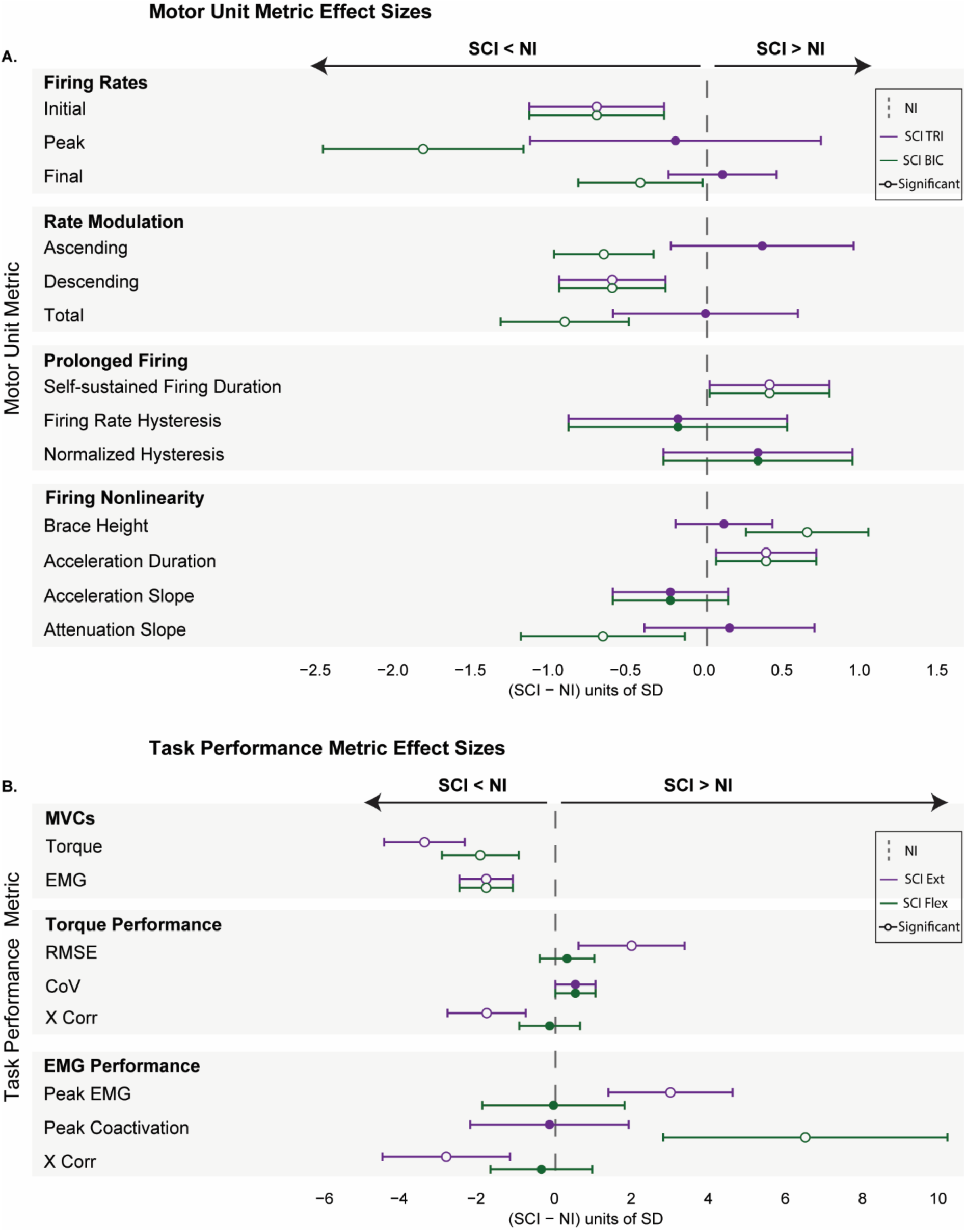
Summary of group contrast effect sizes for all group-level analysis models. (A) Summary of all MU metric models. (B) Summary of MVC and task performance metric models. For each model, the standardized effect size was calculated by dividing the group contrast estimate by the model’s residual standard error. For models using transformed data, effect size is reported on model scale to allow for standardized comparisons. The NI group was used as the reference for each contrast and is represented as the vertical dashed line at 0. Plotted data bars represent the SCI in reference to the NI group and the 95% CI of the contrast (Green = BIC; Purple = TRI). Data that falls to the left of the dashed line indicates the SCI group is lower than the NI group, and data that falls to the right indicates the SCI group is higher than the NI group. Open circles indicate the group contrast is significant (p > 0.05). SCI BIC: n = 20, SCI TRI: n = 15, NI BIC: n = 18, NI TRI: n = 18.

#### Relationships between maximum voluntary torque output and motor unit firing characteristics

##### MVCs

The results from the linear model revealed significant differences in the subgroups for TQ_MVC_. The low-strength SCI subgroup had significantly lower values in flexion (28.0 ± 4.3 Nm) compared to the high-strength SCI (57.6 ± 4.05 Nm; p < 0.001, d = 2.43) and NI (67.2 ± 3.04; p < 0.001, d = 3.2) subgroups. This was also true for extension, with low-strength SCI (17.7 ± 6.6 Nm) having significantly lower values than both high-strength SCI (62.6 ± 7.1 Nm; p < 0.001, d = 2.6) and NI (78.4 ± 4.3 Nm; p < 0.001, d = 3.5).

In support of the cluster analysis, there were no differences between the high-strength SCI and the NI subgroups in TQ_MVC_ for either direction. A similar pattern was observed for EMG during MVC. For BIC during flexion, the low-strength SCI group (0.738 ± 0.21 mV) had significantly lower values compared with both the high-strength SCI (1.55 ± 0.20 mV; p = 0.02, d = 1.4) and NI (2.06 ± 0.15 mV; p < 0.001, d = 2.2) subgroups. However, in TRI during extension, low-strength SCI (0.818 ± 0.28 mV) was only significantly lower than the NI subgroup (1.88 ± 0.19 mV; p = 0.01, d = 1.42), and there was no difference from the high-strength SCI group (0.958 ± 0.30 mV; p = 0.9). There was also a significant difference between high-strength SCI and NI for TRI during extension (p = 0.04, d = 1.2), but not in BIC during flexion (p = 0.1).

##### BIC motor unit firing characteristics and task performance

The linear mixed effects models revealed several significant differences in the rate coding metrics between strength subgroups. For simplicity, only the p-values and effect sizes for significant group differences are reported here, but the full list of estimated marginal means for each outcome can be found in Table 3 and all model contrasts are depicted in figure 8. When compared to the high-strength SCI subgroup, low-strength SCI had significantly lower peak firing rates (p = 0.013, d = 1.13), descending rate modulation (p = 0.038, d = 0.68) and total rate modulation (p = 0.016, d = 0.76). The low-strength SCI also had significantly lower values that the NI subgroup (p < 0.001) for each rate-coding metric (initial firing rate: d = 1.1; peak firing rate: d = 2.2; final firing rate: d = 0.66; ascending rate modulation: d = 0.87; descending rate modulation: d = 1.3; total rate modulation: d = 1.4). Similarly, the high-strength SCI subgroup was significantly lower than the NI subgroup for peak firing rate (p = 0.002, d = 1.1), ascending rate modulation (p = 0.02, d = 0.47), descending rate modulation (p = 0.03, d = 0.63), and total rate modulation (p = 0.03, d = 1.4).

**Figure 8.**
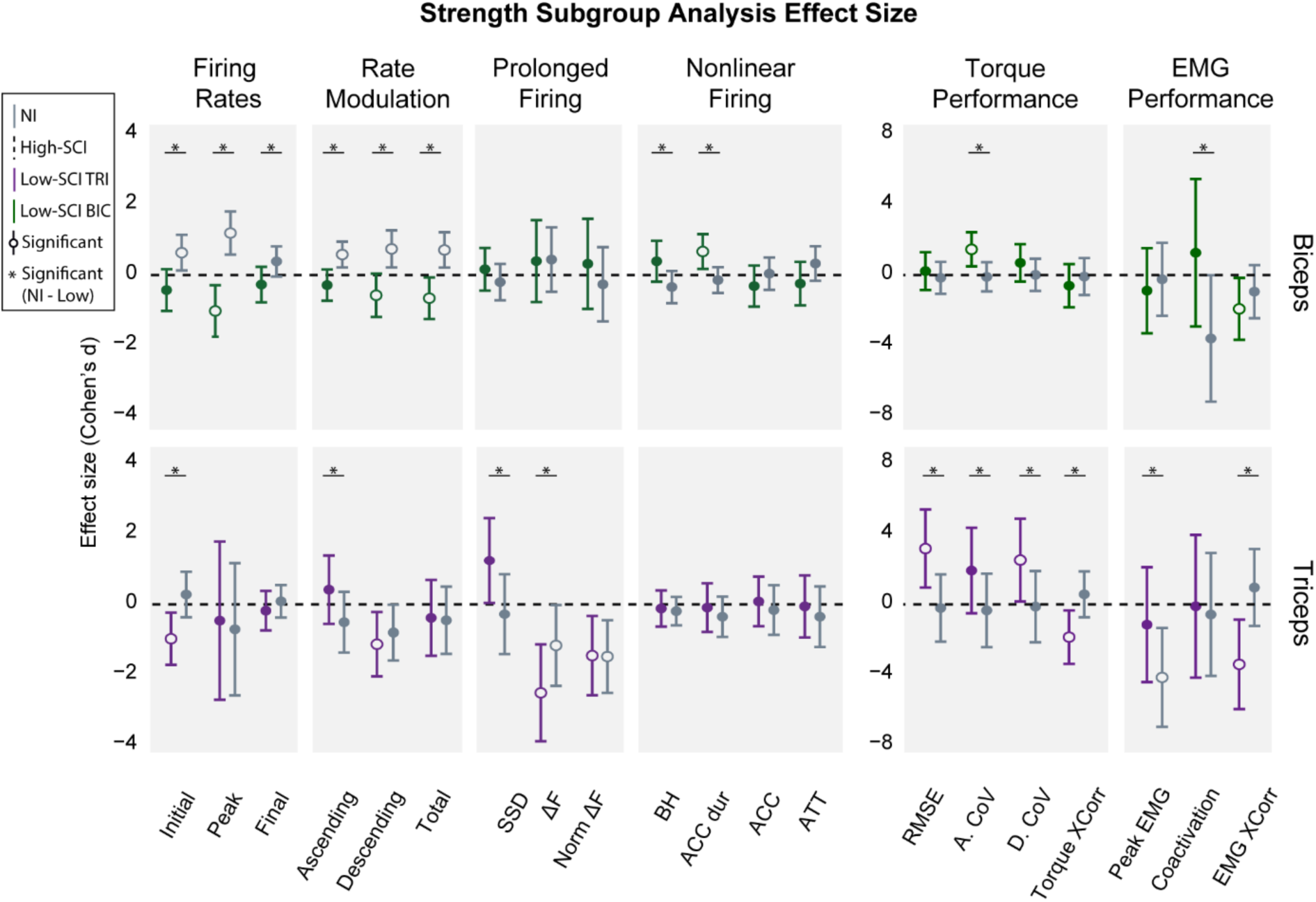
Summary of contrast effect sizes for all strength subgroup-level analysis models. Effect size was calculated as previously described for all 3 contrasts of interest. High-strength SCI was used as the reference group for both muscles and is represented by the black dashed line in each panel. The BIC low-strength SCI data is plotted in green, and the TRI low-strength SCI data is plotted in purple. NI BIC and TRI are both plotted in gray. Data that falls above the dashed line indicates that subgroup has higher values that the high-strength SCI subgroup, and data below indicates that subgroup has lower values compared to the high-strength SCI subgroup. Open circles indicate a group contrast with high-strength SCI is significant. Significance bars with * indicate a significant contrast between low-strength SCI and NI. Abbreviations: SSD, self-sustained duration; BH, brace height; ACC dur, acceleration duration; ACC, acceleration phase; ATT, attenuation phase; RMSE, torque root mean square error; A. CoV, ascending torque coefficient of variation; D. CoV, descending torque coefficient of variation; Torque XCorr, Torque-to-target cross correlation; EMG XCorr, EMG-to-target cross correlation. SCI BIC: n = 20, SCI TRI: n = 15, NI BIC: n = 18, NI TRI = 18.

**Table 3.** Flexion strength subgroup estimated marginal means. ± standard error. Abbreviations: pps, pulses per second; %rTri, percent right triangle; RMSE, root mean square error; CoV, coefficient of variation; MVC, maximal voluntary contraction; Nm, Newton meter; mV, millivolt.

| Outcome | High-strength SCI | Low-strength SCI | NI |
| --- | --- | --- | --- |
| Initial Firing Rate (pps) | 9.36 ± 0.581 | 7.83 ± 0.636 | 10.9 ± 0.453 |
| Peak Firing Rate (pps) | 16.4 ± 0.579 | 13.7 ± 0.652 | 19 ± 0.455 |
| Final Firing Rate (pps) | 7.68 ± 0.433 | 6.72 ± 0.477 | 8.38 ± 0.339 |
| Ascending Rate Modulation (pps) | 6.92 ± 0.431 | 5.76 ± 0.478 | 8.31 ± 0.332 |
| Descending Rate Modulation (pps) | 8.8 ± 0.65 | 6.7 ± 0.71 | 10.8 ± 0.505 |
| Total Rate Modulation (pps) | 9.76 ± 0.54 | 7.7 ± 0.595 | 11.4 ± 0.423 |
| Ascending Dominant (probability) | 0.328 ± 0.047 | 0.382 ± 0.056 | 0.25 ± 0.033 |
| Self-sustained duration (%) | 22.2 ± 6.18 | 23.8 ± 6.73 | 13 ± 4.79 |
| ΔF (pps) | 3.93 ± 0.584 | 4.41 ± 0.769 | 4.48 ± 0.482 |
| Normalized ΔF (a.u.) | 0.379 ± 0.0532 | 0.406 ± 0.0643 | 0.332 ± 0.044 |
| Brace Height (%rTri) | 0.437 ± 0.0263 | 0.478 ± 0.0332 | 0.373 ± 0.021 |
| Attenuation Slope (pps/% MVC) | 0.422 ± 0.0759 | 0.287 ± 0.0939 | 0.506 ± 0.059 |
| Acceleration Slope (pps/% MVC) | 2.75 ± 0.305 | 2.05 ± 0.386 | 2.64 ± 0.238 |
| Acceleration Phase Duration (sec) | 1.28 ± 0.0898 | 1.61 ± 0.119 | 1.13 ± 0.0687 |
| Torque MVC (Nm) | 57.6 ± 4.05 | 28 ± 4.3 | 67.2 ± 3.04 |
| EMG MVC (mV) | 1.55 ± 0.198 | 0.738 ± 0.210 | 2.06 ± 0.148 |
| Torque RMSE (%) | 0.576 ± 0.0857 | 0.627 ± 0.0929 | 0.539 ± 0.065 |
| Torque Ascending CoV (%) | 3.26 ± 0.29 | 4.51 ± 0.308 | 3.17 ± 0.218 |
| Torque Descending CoV (%) | 4.19 ± 0.615 | 5.33 ± 0.652 | 4.21 ± 0.461 |
| Torque-to-Target (Pearson r) | 0.995 ± 0.001 | 0.994 ± 0.001 | 0.995 ± 0.001 |
| Peak EMG (% EMG MVC) | 20.8 ± 3.02 | 17.6 ± 3.2 | 19.9 ± 2.26 |
| Muscle Coactivation | 0.427 ± 0.0729 | 0.49 ± 0.0773 | 0.247 ± 0.055 |
| EMG-to-Target (Pearson r) | 0.978 ± 0.004 | 0.959 ± 0.009 | 0.971 ± 0.005 |

There were no significant differences between any of the BIC subgroups for the prolonged firing rate metrics. However, for non-linear firing rate metrics, the low-strength SCI had significantly longer acceleration phase durations compared to the high-strength SCI (p = 0.038, d = 0.56) and the NI subgroup (p = 0.002, d = 0.81). Brace height was also significantly increased in the low-strength SCI compared to the NI subgroup (p = 0.01, d = 0.73). For task performance, there were no significant differences between the high-strength SCI and NI subgroups. However, the high-strength SCI had significantly lower ascending torque CoV (p = 0.006, d = 1.5) and significantly greater EMG-to-Target correlation values (p = 0.04, d = 1.9) compared to the low-strength SCI subgroup. The low-strength SCI group also significantly differed from the NI subgroup, with higher ascending torque CoV (p = 0.001, d = 1.6) and muscle coactivation (p = 0.02, d = 4.7).

**Table 3** Flexion strength subgroup estimated marginal means ± standard error. Abbreviations: pps, pulses per second; %rTri, percent right triangle; RMSE, root mean square error; CoV, coefficient of variation; MVC, maximal voluntary contraction; Nm, Newton meter; mV, millivolt.

##### Triceps motor unit firing characteristics and task performance

As with the previous section, the p-values and effect sizes are listed for significant group contrasts, but the full comparison of effect sizes can be found in figure 8 and estimated means are listed in Table 4. The linear mixed effects models revealed a different pattern of group differences in the TRI than what we observed in the BIC, with only a few significant differences in rate-coding metrics. The SCI low-strength subgroup had significantly lower initial firing rates compared to both the high-strength SCI (p = 0.008, d = 1.1) and NI (p < 0.001, d = 1.3) subgroups. The low-strength SCI also had significantly greater ascending rate modulation compared to the NI subgroup (p = 0.03, d = 0.9), and significantly lower descending rate modulation compared to the high-strength SCI. In line with these results, the low-strength SCI had a significantly higher probability of having ascending dominant rate modulation compared to both the high-strength SCI (Low/High odds ratio = 2.33 ± 0.81, p = 0.039) and the NI (Low/NI odds ratio = 2.99 ± 0.86, p < 0.001). This scenario is demonstrated in figures 9H and 10, where TRI MUs from low-strength SCI individuals show greater ascending rate modulation compared to descending rate modulation. There were no additional differences in rate coding metrics. There were, however, significant differences in prolonged firing and onset-offset hysteresis. The low-strength SCI subgroup had significantly greater self-sustained duration compared to the NI subgroup (p = 0.01, d = 1.5). The high-strength SCI had significantly higher ΔF values compared to both the low-strength SCI (p = 0.001, d = 2.6) and NI (p = 0.04, d = 1.3) subgroups. The low-strength SCI also had significantly lower ΔF values than the NI subgroup (p = 0.03, d = 1.3). For normalized ΔF, the high-strength SCI group remained significantly higher than both low-strength SCI (p = 0.01, d= 1.6) and NI (p = 0.006, d = 1.6) subgroups, while the low-strength SCI and NI subgroup values were similar. We did not observe any significant differences in the non-linear firing rate metrics for the triceps.

**Figure 9.**
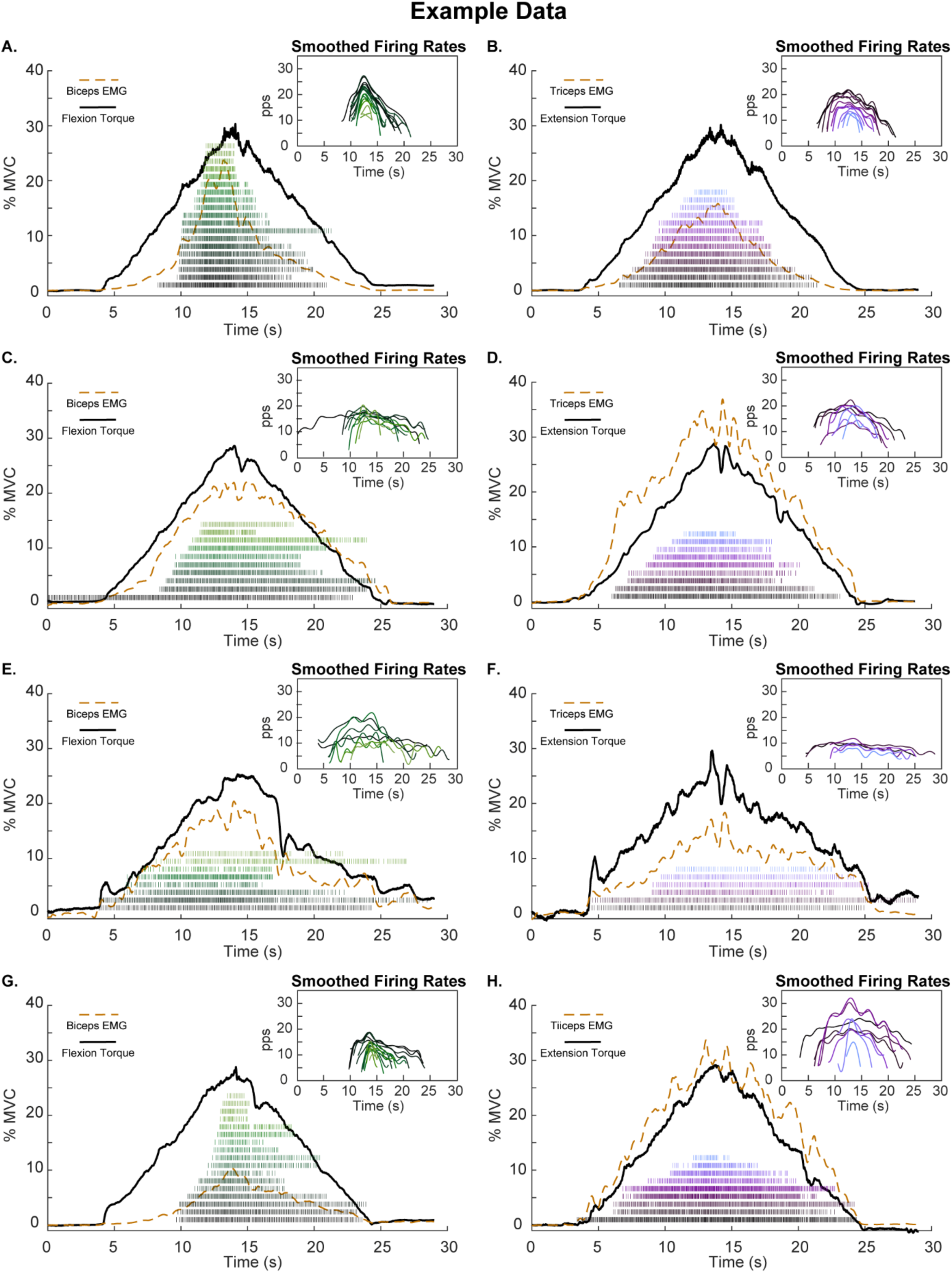
Example data from 4 participants. (A-B) Flexion and extension trials, respectively, from NI participant NI0C. (C-D) Flexion and extension trials, respectively, from high-strength BIC and high-strength TRI SCI participant SCI20. (E-F) (A-B) Flexion and extension trials, respectively, from low-strength BIC and low-strength TRI SCI participant SCI11. (G-H) (A-B) Flexion and extension trials, respectively, from SCI participant with high-strength BIC and low-strength TRI (SCI0C). Each plot includes the isometric contraction torque trace (black), the agonist EMG envelope (orange dashed line), raster plot of instantaneous MU discharge times to show recruitment distribution and MU yield, and the smoothed MU firing rates for firing rate modulation comparison. SCI BIC: n = 3, SCI TRI: n = 3, NI BIC n = 1, NI TRI n = 1.

**Table 4.** Extension strength subgroup estimated marginal means. ± standard error. Abbreviations: pps, pulses per second; %rTri, percent right triangle; RMSE, root mean square error; CoV, coefficient of variation; MVC, maximal voluntary contraction; Nm, Newton meter; mV, millivolt.

| Outcome | High-strength SCI | Low-strength SCI | NI |
| --- | --- | --- | --- |
| Initial Firing Rate (pps) | $9 \pm 0.649$ | $6.48 \pm 0.597$ | $9.4 \pm 0.401$ |
| Peak Firing Rate (pps) | $17.1 \pm 1.03$ | $16.4 \pm 0.983$ | $16 \pm 0.628$ |
| Final Firing Rate (pps) | $6.9 \pm 0.466$ | $6.22 \pm 0.47$ | $6.84 \pm 0.289$ |
| Ascending Rate Modulation (pps) | $8.3 \pm 0.868$ | $9.04 \pm 0.799$ | $6.82 \pm 0.534$ |
| Descending Rate Modulation (pps) | 9.99 ± 0.836 | 9.8 ± 0.808 | 9.45 ± 0.505 |
| Total Rate Modulation (pps) | 11.3 ± 0.895 | 10.2 ± 0.822 | 10.1 ± 0.549 |
| Ascending Dominant (probability) | 0.281 ± 0.053 | 0.476 ± 0.057 | 0.233 ± 0.304 |
| Self-sustained duration (%) | 9.94 ± 6.75 | 27.59 ± 6.08 | 5.76 ± 6.75 |
| ΔF (pps) | 5.63 ± 0.592 | 2.49 ± 0.597 | 4.1 ± 0.376 |
| Normalized ΔF | 0.523 ± 0.0438 | 0.351 ± 0.0447 | 0.348 ± 0.029 |
| Brace Height (%rTri) | 0.519 ± 0.0211 | 0.492 ± 0.0238 | 0.482 ± 0.013 |
| Attenuation Slope (pps/% MVC) | 0.382 ± 0.0909 | 0.338 ± 0.0854 | 0.26 ± 0.0576 |
| Acceleration Slope (pps/% MVC) | 2.89 ± 0.534 | 2.84 ± 0.498 | 2.37 ± 0.336 |
| Acceleration Phase Duration (sec) | 1.79 ± 0.157 | 1.66 ± 0.162 | 1.5 ± 0.0985 |
| Torque MVC (Nm) | 62.6 ± 7.08 | 17.7 ± 6.55 | 78.4 ± 4.33 |
| EMG MVC (mV) | 0.958 ± 0.303 | 0.818 ± 0.281 | 1.880 ± 0.186 |
| Torque RMSE (%) | 0.474 ± 0.195 | 1.22 ± 0.18 | 0.425 ± 0.119 |
| Torque Ascending CoV (%) | 3.79 ± 1.45 | 6.86 ± 1.34 | 3.24 ± 0.889 |
| Torque Descending CoV (%) | 4.37 ± 1.32 | 8.14 ± 1.22 | 4.18 ± 0.808 |
| Torque-to-Target (Pearson r) | 0.995 ± 0.001 | 0.991 ± 0.002 | 0.996 ± 0.0004 |
| Peak EMG (% EMG MVC) | 27.8 ± 3.01 | 24.9 ± 2.79 | 17.5 ± 1.84 |
| Muscle Coactivation | 0.168 ± 0.0478 | 0.164 ± 0.0442 | 0.15 ± 0.0293 |
| EMG-to-Target (Pearson r) | 0.985 ± 0.004 | 0.962 ± 0.009 | 0.989 ± 0.002 |

For task performance, when compared to the high-strength SCIs, the low-strength SCIs had significantly higher torque RMSE (p = 0.009, d = 3.2) and descending torque CoV (p = 0.045, d = 2.5), and lower Torque-to-Target cross correlation (p = 0.02, d = 1.9) and EMG-to-Target cross correlation (p = 0.01, d = 3.4). When compared to the NI subgroup, the low-strength SCI again had significantly higher RMSE and CoV values (Torque RMSE: p < 0.001, d = 3.4; Ascending Torque CoV: p = 0.03, d = 2.3; Descending Torque CoV: p = 0.01, d = 2.6) and lower Torque-to-Target cross correlation (p < 0.001, d = 2.4) and EMG-to-Target cross correlation (p <0.001, d = 4.4). For Peak agonist EMG amplitude during the contraction, the NI subgroup had significantly lower values compared to both the high-strength SCI (p = 0.007, d = 4.1) and low-strength SCI (p = 0.04, d = 2.9) subgroups, but there was no difference between SCI subgroups for this metric.

## Discussion

The results from this study demonstrate muscle-specific differences in MU firing characteristics following cervical level spinal cord injury. The group-level analysis revealed that BIC MUs in the SCI group displayed a significant reduction in rate-coding behaviors compared with the NI group. This was accompanied by significant group differences in ascending firing rate non-linearity (i.e. Brace Height and attenuation slope). Differences in rate coding behaviors were not observed in the group-level analysis of the TRI, but the strength subgroup-level analysis of the triceps revealed significant differences in PIC-related metrics that promote prolonged firing (i.e. self-sustained duration, ΔF) between the high-strength SCI and low-strength SCI. Such relationships indicate a possible link between motoneuron properties, and their inputs, and TRI strength that may have been masked when grouping the heterogeneous participants together.

### Muscle-specific differences in rate-coding behavior

The group-level analysis revealed more prominent differences in rate-coding for the BIC compared to the TRI. Initial and peak firing rates, as well as rate modulation (ascending, descending, and total), were all significantly reduced in the BIC of the SCI group when compared to the NI group. For TRI, group differences were revealed only when averaged across both muscles for final firing rate and descending rate modulation. These muscle-specific differences after SCI are in line with what has been reported previously by Wiegner *et al*. (1993), where firing rates of BIC were significantly reduced compared to control participants as opposed to TRI firing rates after SCI that were similar to control participants. This could be attributed to muscle-specific changes in afterhyperpolarization (AHP), a predominant mechanism that has been linked to reduced firing rates (Wienecke, *et al.,* 2009). Although this has not yet been investigated in the upper limb, Duan and colleagues (2025) have recently reported an increase in AHP duration in the tibialis anterior (TA), but not in soleus or medial gastrocnemius, of individuals with SCI compared to non-injured controls. They suggest this differential increase could be attributed to differences in corticospinal input, and because the TA receives more corticospinal projections than the triceps surae, these MUs may be more sensitive to reductions in this input following an injury. Kizyte and colleagues (2025) provide further evidence for this link of increased AHP duration and reduced firing rate after SCI in their study that found SCI participants to have significantly lower firing rates in the TA compared to non-injured controls, but they found no group differences in soleus firing rates. Since significant differences were observed in the flexors, but not extensors of the leg, it is plausible that similar flexor-extensor differences in the AHP exist in the arms between the BIC (i.e., flexors) and TRI (i.e., extensors), but further work is required to substantiate this suggestion.

An alternative explanation is that there are muscle-specific patterns of excitation-inhibition coupling, of which several schemes have been proposed. In a push-pull scheme, as excitatory drive to the MU increases, there is an opposite decrease in inhibition which would facilitate MU firing rate modulation. Similarly, a constant level of background inhibition during increasing and decreasing excitation also allows for firing rate modulation, albeit to a lesser extent. With a proportional scheme, however, where excitation and inhibition increase and decrease in parallel, there is reduced net excitatory input to the MUs and attenuated firing rate modulation (Powers *et al.,* 2012). Changes in presynaptic inhibition (Mailis & Ashby, 1990; Faist *et al.,* 1994; Aymard *et al.,* 2000), post-activation depression (Schindler-Ivens & Shields, 2000; Nito *et al.,* 2026), and reciprocal inhibition (Okuma *et al.,* 2002; Crone *et al.,* 2003) are commonly reported after SCI and may contribute to altered patterns of excitation-inhibition coupling during voluntary force output.

Although we did not explicitly test differences in reciprocal inhibition, we observed a significant increase in antagonist muscle coactivation (i.e., triceps) during flexion in the SCI group compared to the NI group. This was similarly observed in previous studies (Thomas *et al.,* 1997; Cremoux *et al.,* 2016), and it is likely that the increased TRI activity during flexion imposed reciprocal inhibition onto the BIC MUs, possibly contributing to the dampened rate coding metrics we observed (Gomes *et al.,* 2024). This suggestion is supported by the lack of group differences in antagonist muscle coactivation (i.e. BIC) during extension and the preserved peak firing rates of the TRI in the SCI group. Although previous studies have shown there to be no difference in the amount of Ia reciprocal inhibition imposed between the BIC and TRI in the neurologically intact population (Katz *et al.,* 1991), we are unaware of any comparisons made after SCI. Although further investigation is needed, given the post-SCI changes observed in other muscle groups (Boorman *et al.,* 1996; Xia & Rymer, 2005) and increase in reciprocal facilitation with greater motor impairment (Okuma *et al.,* 2002), it is plausible that a similar imbalance exists between the BIC and TRI.

### Muscle-specific differences in non-linear and prolonged firing behavior

Decades of work in reduced preparations have revealed the critical role of PICs in modulating motoneuron sensitivity to excitatory and inhibitory synaptic inputs (Delgado-Lezama *et al.,* 1997; Svirskis & Hounsgaard, 1998; Lee & Heckman, 2000; Hultborn, *et al.,* 2003). In a neurologically intact system, PICs are facilitated by the brainstem derived monoamines 5HT and NA, but following SCI, there can be a significant reduction in the amount of intact bulbospinal projections below the level of injury, limiting PIC facilitation and making it difficult for full activation of MUs. This in turn reduces muscle force production, which can limit the individual’s ability to perform functional movements. Indeed, it has been demonstrated that motoneuron excitability is significantly reduced in the acute and sub-acute phases of SCI, during a period referred to as spinal shock. However, moving into the chronic phase of the injury there is a recovery of excitability that is often accompanied by the manifestation of involuntary muscle activity or spasticity (Hiersemenzel, *et al.,* 2000; Ditunno *et al.,* 2004).

The mechanisms underlying this restoration of excitability is the upregulation of constitutively active 5HT2c and NAα_1_ receptors that facilitate PICs in the absence of endogenous monoamines. This has been demonstrated in both the motor complete (Murray *et al.,* 2010; 2011) and motor incomplete (Tysseling, *et al.,* 2017) rodent models of SCI. D’Amico and colleagues (2013) have also explored this upregulation in humans, where oral administration of cyproheptadine, an inverse agonist to 5-HT2 and NAα1 receptors, reduced the PIC-mediated long-lasting reflex in individuals with motor complete or incomplete injuries.

Due to this reported upregulation, we investigated whether an increase in non-linear and prolonged MU firing behavior in the SCI group is a mechanism underlying augmented muscle strength and whether muscle-specific changes may enhance recovery in one muscle versus another. The results of our group-level analysis revealed that brace height was significantly increased in the BIC of the SCI group compared to the NI group. Because this is a measure of the non-linearity in ascending firing rate caused by slow PIC activation near the onset of firing, larger values indicate more neuromodulation (Beauchamp, *et al.,* 2023). Conversely, attenuation slopes in the BIC were significantly lower in the SCI group compared to the NI group. This is similar to what was seen in the rate-coding metrics and again suggests a possible either a shift in the excitation-inhibition coupling after SCI or limited excitatory input to motoneuron pools. Computer simulations have shown that the attenuation slope is largely unaffected by changes in neuromodulation but decreases when there is a shift from a reciprocal (or push-pull) to a proportional excitation-inhibition pattern (Beauchamp *et al.,* 2023). In this scenario, the parallel increase in excitation and inhibition reduces the amount of net excitation and suppresses firing rate modulation.

Also referred to as the secondary firing rate, the acceleration slope is quantified as the initial steep increase in firing rate following recruitment, which is driven by PIC activation and synaptic amplification. This phase typically lasts for less than 2 seconds before transitioning to the slower tertiary firing rate (i.e., the attenuation slope) as the PIC becomes fully activated and increases in MU firing rate become attenuated. In the SCI group, the acceleration durations were longer in both muscles, potentially indicating reduced subthreshold contributions of PICs. An alternative explanation could be that secondary firing range durations are longer due to slower kinetics of the voltage sensitive L-type Ca^2+^ and Na^+^ channels, or that there is a reduced contribution of fast-activation of Na^+^ and shift to a greater reliance on slower L-type Ca^2+^ channels to accelerate motoneuron firing. Furthermore, simulations indicate that the acceleration slope is less sensitive to changes in levels of neuromodulation and inhibition patterns than brace height. This is potentially due to the normalization procedures applied to brace height that reduce variability, in comparison to the use of raw acceleration slope values (Beauchamp, *et al.,* 2023). Therefore, the lack of group differences here is unsurprising.

The results of our group-level analysis for firing rate hysteresis (ΔF) did not reveal significant differences between the SCI and NI groups for either BIC or TRI. We also did not observe differences between muscles for either group, which contrasts previous work by Wilson and colleagues (2015). Their work revealed greater ΔF in the TRI compared with BIC PICs in neurologically intact participants. The lack of muscle-specific differences in ΔF in our current study may be due to methodological differences. Specifically, the tested position in our study resulted in elongated TRI and shortened BIC muscle lengths. Muscle length has been shown to modulate ΔF values, with longer lengths showing reduced values compared to short lengths (Beauchamp *et al.,* 2025; Goreau *et al.,* 2025). Muscle-specific differences may also be affected by a difference in contraction amplitudes. The previous study only tested MU firing during contractions to 10% of their maximum voluntary torque, and ΔF has also been shown to change with contraction intensity (Škarabot *et al.,* 2025).

Although we did not observe group difference in ΔF, there was a significant increase in self-sustained MU firing duration in both muscles, indicating increased motoneuron excitability and a possible shift in excitation-inhibition that promotes sustained firing. Computer simulations have demonstrated the sensitivity of firing rate hysteresis to changes in both neuromodulation and inhibition. Although increases in neuromodulation increase the value of ΔF, it becomes less pronounced with a strong proportional excitation-inhibition scheme (Chardon *et al.,* 2024). This may partially explain why there were no significant differences observed in firing rate hysteresis for the group-level analysis in this study despite the observed changes in brace height in the BIC and self-sustained firing duration in both BIC and TRI. However, the impact of different excitation-inhibition patterns on self-sustained firing durations has not yet been explored to our knowledge.

### Functionallimitations in performance that introduce variability in MU behaviors

Despite the significant group differences observed in the BIC, it is important to note the broad range of strength we observed in our SCI participants for both flexion and extension. Several of the participants produced torque and muscle activity during MVCs within the range of the NI group (refer to fig. 3), with their MU firing profiles indistinguishable from that of a non-injured participant. On the opposite end of that spectrum, many of the impaired individuals, who produced much lower TQ_MVC_ values, produced MU firing patterns with varied amounts of rate modulation and sustained MU firing, in some cases even while they were supposed to be at rest. The variety in firing behaviors can be appreciated by comparing the examples provided in figure 9, which depicts BIC (left column) and TRI (right column) data from four example participants with varying degrees of impairment.

The top row (Fig 9A-B) depicts trial examples from a male NI participant (NI06: Flexion TQ_MVC_ = 92.4 Nm; Extension TQ_MVC_ = 93.9 Nm). The second row (Fig 9C-D) shows data from a male SCI with high-strength BIC and high-strength TRI (SCI20: AIS-C, LOI – C7; Flexion TQ_MVC_ = 58.30 Nm; Extension TQ_MVC_ = 62.04 Nm). Compared to the NI, there is similar, but slightly reduced, rate modulation in the BIC, with a tonically active MU at the start of the trial. The third row (Fig 9D-E) shows data from one of the more-impaired male SCI participants with low-strength BIC and low-strength TRI (SCI11: AIS-C, LOI-C4; Flexion TQ_MVC_ = 17.38 Nm; Extension TQ_MVC_ = 9.9 Nm). Their lower level of functional control is reflected in the higher error and CoV observed in the torque profiles. Although there is some MU rate modulation in the BIC (Fig 9D), the firing rates are much more variable and disordered compared to the first two participants. Rate modulation deficits are even more visible in this individual’s TRI (Fig 9E), and there are several MUs in both muscles exhibiting strong sustained firing throughout the duration of the trial. This behavior was absent in the NI group but was commonly observed in the SCI group, and particularly in the low-strength SCIs, with 13 of the 20 participants producing trials where 1-6 MUs sustained their activity beyond the torque trace. The bottom row (Fig 9G-H) depicts data from one of the 3 male SCI participants that was classified as high-strength BIC but low-strength TRI (SCI06: AIS-A, LOI-C6; Flexion TQ_MVC_ = 47.3 Nm; Extension TQ_MVC_ = 22.44 Nm). Here, the BIC again shows rate modulation, but at a more condensed range compared to the example from NI06. However, in the TRI there is a higher amount of rate modulation compared to NI06, and there also appears to be a shift away from the typical “onion-skin” firing rate scheme (Pearcey & Rymer, 2022) that is clearly shown in 9B, where earlier recruited MUs tend to have higher peak firing rates and larger rate modulation ranges. Although there are fewer decomposed MUs, figure 9H shows a much less organized scheme in comparison.

### Strength-related subgroup analysis

To better explore this observed variability, we created strength-related subgroups using a clustering analysis of TQ_MVC_ values. The subgroup analysis of the rate-coding behaviors in the BIC showed the same pattern we observed in the group-level analysis. However, the suppression of these metrics is exemplified in the low-strength SCI subgroup compared to the high-strength SCI subgroup. These results suggest that the BIC are more susceptible to dampened rate modulation following SCI, and that recovery or preservation of typical rate-coding behaviors is either a prerequisite or may promote strength in this muscle. Due to the reliance on excitatory synaptic input, residual descending projections or an upregulation in excitatory reticulospinal ionotropic input (Sangari & Perez, 2020) following an injury, may facilitate MU firing rate modulation in the BIC of the higher strength individuals. There were fewer subgroup-level differences in rate-coding metrics for the TRI, although some key differences were revealed. The weaker-strength SCI subgroup had elevated ascending rate modulation compared to the NI subgroup, but lesser descending rate modulation compared to the high-strength SCI subgroup. This is reflected in their significantly lower initial firing rates and increased probability of having a greater proportion of ascending vs. descending rate modulation.

The rate-coding results in the TRI also correspond to the significant differences we observed in prolonged firing rates. Here, the low-strength SCI subgroup had significantly more self-sustained firing compared to the NI subgroup, indicating an increase in PICs (i.e. excitability). It may be the case that altered inhibition patterns prevent rate modulation during the descending phase of the contraction and the MUs get stuck in a heightened state of excitability (i.e., stuck on a plateau potential). Limited descending rate modulation in the low-strength subgroup would also result in lower ΔF values, which is what we observed. When ΔF is normalized to rate modulation, there is no difference between the low-strength SCI and NI subgroups, but the high-strength SCI do have significantly greater values. As previously discussed in the group-level analysis section, this observed difference in ΔF between the high-strength and the low-strength SCI could be due to differences in inhibition (Chardon *et al.,* 2024).

For the BIC subgroup analysis, the only observed difference in non-linear metrics between the high- and low-strength SCI subgroups occurred in acceleration phase duration, where low-strength SCI had significantly longer durations than high-strength. Low-strength SCI also had significantly greater brace height values compared to the NI group. This indicates that the results from the overall group-level analysis for these metrics may be driven by weaker individuals with SCI and could be linked to reduced functional recovery. The discrepancy between these subgroups for both BIC and TRI highlights the need to be cautious when grouping together individuals with heterogenous injuries that result in diverse levels of strength and functionality. In doing so, there is the potential risk of masking significant findings that could inform rehabilitation strategies.

### Important strength and task performance considerations

TQ_MVC_ and EMG_MVC_ were reduced in the SCI group during both flexion and extension. In the NI group, extension torque exceeded flexion torque, whereas no directional difference was observed in the SCI group. This pattern suggests a greater loss of elbow extension strength relative to flexion after SCI, consistent with prior reports of asymmetric recovery between the BIC and TRI (Ditunno et al., 1992; Calancie et al., 2004; McKay et al., 2011; Balbinot et al., 2023). However, this interpretation should be made cautiously, as the greater extension torque in the NI group may also reflect differences in trunk stabilization in the test position. Nevertheless, the positive relationship between torque and muscle activity across participants indicates that TQ_MVC_ remains a reasonable index of elbow flexion and extension strength.

During the 30% TQ_MVC_ ramp contractions, the SCI group showed greater errors in torque during extension, with higher RMSE and lower torque-to-target cross-correlation values than the NI group. In contrast, there were no group differences in these metrics during flexion. Although torque variability was higher in the SCI group for both flexion and extension, this difference did not reach significance. The improved task performance during flexion may be a combination of increased antagonist muscle coactivation and reduction in peak firing rates. In contrast, BIC coactivation during extension was generally low but peak firing rates in the TRI were often quite large, even in the more impaired individuals. This likely contributed to less joint stability and target accuracy of the movement. Agonist EMG showed a similar pattern, with greater variability in TRI envelope shape and amplitude during extension. In contrast, no significant group differences were observed in these metrics for the BIC during flexion.

As previously discussed, antagonist muscle coactivation was higher during flexion in the SCI group, while coactivation during extension was similar between groups. One limitation of this analysis is that MVC normalization may have inflated values in participants with greater impairment, especially with the four SCI participants who could not complete the extension ramp and were thus excluded from the triceps MU analysis. These individuals generated measurable TRI EMG during extension MVCs, but the peak values were lower than what was recorded during their flexion MVCs. For these participants, their normalized antagonist activity appears to dominate the ramp contraction. This is not a unique phenomenon and has been previously reported in SCI individuals exhibiting functional strength imbalances between BIC and TRI (Cremoux *et al.,* 2016). We explored using the TRI peak EMG recorded during their flexion MVC as the normalization reference. Although this reduced the coactivation values, it did not change the overall group results, and the same was true when these participants were excluded entirely from the coactivation analysis. We therefore retained the true extension MVC values because they best represented maximal TRI EMG during joint torque isolation.

### Further considerations

An important limitation in our study, which is often overlooked when assessing MU firing characteristics in people with motor impairment, is variability in the Peak EMG magnitudes and overall EMG envelope shapes across participants despite the consistent magnitude and shape of the torque profiles. This limitation, however, was not isolated to the SCI group, as there was also variability in the NI group’s EMG properties. This highlights two important considerations: 1) accurate interpretation of all our MU metrics across participants relies on normalizing the magnitude of neural drive to a percentage of MVC; and 2) interpretation of the non-linear MU metrics (i.e. ΔF, Brace Height, etc.) assumes a linear and symmetrical increase and decrease in neural drive. To control for this, we explored conservatively filtering the data to only include trials with similar EMG envelope characteristics. We used the median value of Peak agonist EMG across all trials (18.85% EMG_MVC_), calculated the median absolute deviation of each trial (MAD = 5.02), and set the trial inclusion criteria range as median ± 2 MAD (8.8 – 28.9%). We then filtered out trials with EMG-to-target cross correlation (r) values less than 0.95. Although we did observe a minor change in the magnitude of the group contrast for normalized firing rate hysteresis, it still did not reach statistical significance, and no other notable differences were present in the group analysis. Imposing stricter filtering criteria critically limited the number of participants, rendering it difficult to pursue further. We chose, instead, to report the MU recruitment thresholds used in the LMER models in terms of % EMG_MVC_ and included Peak EMG and EMG-to-Target cross correlation values as covariates when physiologically relevant to account for the variability across participants and trials.

Another aspect of the dataset that requires consideration is the variability in the number of decomposed MUs across participants. Although there were no significant differences in the average number of MUs identified per trial between groups or muscles, there was a large amount of variability across participants within both groups. Furthermore, as previously mentioned, we were unable to calculate the firing rate hysteresis or brace height variables for all the decomposed MUs due to important inclusion criteria within the calculation of these metrics. This is a particular limitation for the SCI group where the number of MUs identified tended to be slightly higher in stronger individuals, and impaired MU behaviors were less likely to meet the criteria for ΔF and brace height calculations. Example data of such a situation is provided in figure 10 (SCI02, AIS-C, LOI-C5; Extension Torque MVC = 2.86 Nm). This is a clear example of exaggerated self-sustained firing in low-strength TRI, where neither ΔF nor the non-linear metrics (i.e., Brace height, etc.) could be calculated for any of the MUs. Lastly, as we have already mentioned, four of the participants with SCI were too impaired to adequately perform the elbow extension task and their trials had to be excluded from the final analysis. Thus, several of the most impaired individuals were not fully represented in the data.

**Figure 10.**
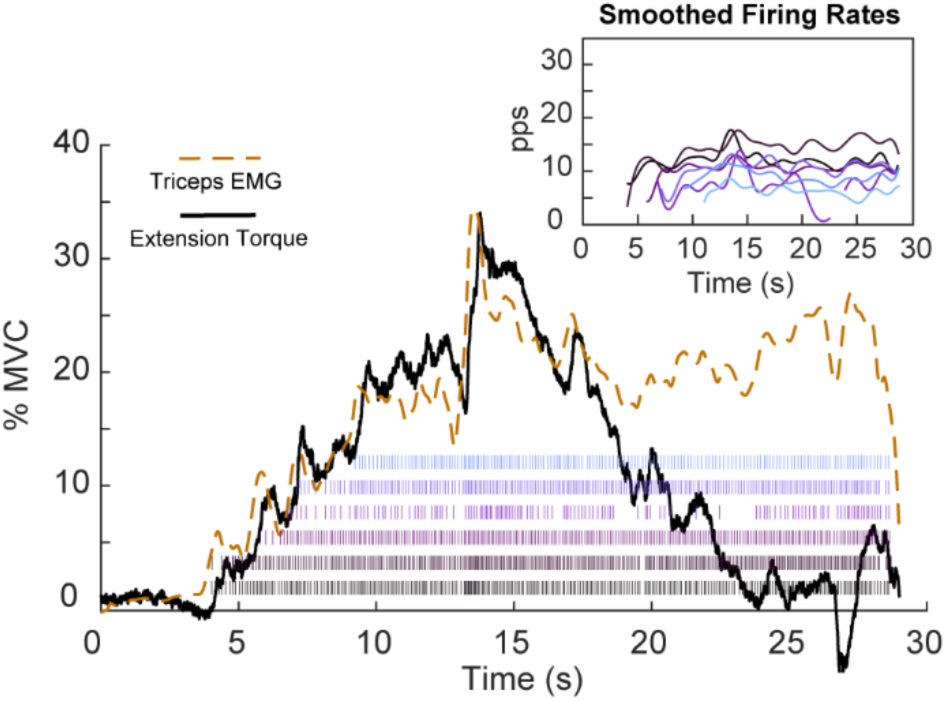
Example data from the triceps of one female SCI participant. AIS impairment level is C and level of injury is C5. Extension torque MVC value was low enough to be classified as low-strength in the strength sub-group analysis.

## CONCLUSION

In this study, we demonstrate substantial muscle-specific differences in upper limb MU firing behavior following cervical level spinal cord injury. Furthermore, we identify distinct MU firing patterns that indicate an increase in neuromodulation and/or a shift in excitation-inhibition coupling that may be linked to preserved or recovered strength after injury. Rehabilitation interventions that promote an increase in motoneuron excitability and restoration of typical inhibition patterns may enhance functional outcomes while suppressing maladaptive behaviors such as self-sustained firing. We also acknowledge and highlight the need for a nuanced approach when analyzing these behaviors in a heterogeneously impaired population that exhibits extensive functional and strength variability.

## REFERENCES

Afsharipour, B., Manzur, N., Duchcherer, J., Fenrich, K. F., Thompson, C. K., Negro, F., … & Gorassini, M. A. (2020). Estimation of self-sustained activity produced by persistent inward currents using firing rate profiles of multiple MUs in humans. Journal of Neurophysiology.

Balbinot, G., Li, G., Kalsi-Ryan, S., Abel, R., Maier, D., Kalke, Y. B., … C Zariffa, J. (2023). Segmental motor recovery after cervical spinal cord injury relates to density and integrity of corticospinal tract projections. Nature communications, 14(1), 723.

Bao, S., & Lei, Y. (2024). Motor unit activity and synaptic inputs to motoneurons in the caudal part of the injured spinal cord. Journal of Neurophysiology, 131(2), 187–197.

Beauchamp, J. A., Khurram, O. U., Dewald, J. P., Heckman, C. J., & Pearcey, G. E. (2022). A computational approach for generating continuous estimates of MU discharge rates and visualizing population discharge characteristics. Journal of Neural Engineering, 19(1), 016007.

Beauchamp, J. A., Pearcey, G. E., Khurram, O. U., Chardon, M., Wang, Y. C., Powers, R. K., … & Heckman, C. J. (2023). A geometric approach to quantifying the neuromodulatory effects of persistent inward currents on individual MU discharge patterns. Journal of Neural Engineering, 20(1), 016034.

Beauchamp, J. A., Pearcey, G. E., Khurram, O. U., Negro, F., Dewald, J. P., & Heckman, C. J. (2025). Intrinsic properties of spinal motoneurons degrade ankle torque control in humans. The Journal of Physiology, 603(8), 2443–2463.

Bennett, D. J., Li, Y., & Siu, M. (2001). Plateau potentials in sacrocaudal motoneurons of chronic spinal rats, recorded in vitro. Journal of neurophysiology, 86(4), 1955–1971.

Binder MD, Powers RK, Heckman CJ (2020) Non-linear Input-Output Functions of Motoneurons. Physiol Bethesda 35:31–39.

Butler, C. L., Sangari, S., Chen, B., & Perez, M. A. (2024). Enhanced inhibitory input to triceps brachii in humans with spinal cord injury. The Journal of physiology, 602(24), 6909–6923.

Calancie, B., Molano, M. R., & Broton, J. G. (2004). EMG for assessing the recovery of voluntary movement after acute spinal cord injury in man. Clinical neurophysiology, 115(8), 1748–1759.

Chardon, M. K., Wang, Y. C., Garcia, M., Besler, E., Beauchamp, J. A., D’Mello, M., … & Heckman, C. J. (2024). Supercomputer framework for reverse engineering firing patterns of neuron populations to identify their synaptic inputs. elife, 12, RP90624.

D’Amico, J. M., Murray, K. C., Li, Y., Chan, K. M., Finlay, M. G., Bennett, D. J., et al. (2013). Constitutively active 5-HT2/alpha1 receptors facilitate muscle spasms after human spinal cord injury. J. Neurophysiol. 109, 1473–1484. doi: 10.1152/jn.00821.2012.

Delgado-Lezama, R., Perrier, J. F., Nedergaard, S., Svirskis, G., & Hounsgaard, J. (1997). Metabotropic synaptic regulation of intrinsic response properties of turtle spinal motoneurones. The Journal of physiology, 504(Pt 1), 97.

Del Vecchio, A., Casolo, A., Negro, F., Scorcelletti, M., Bazzucchi, I., Enoka, R., … C Farina, D. (2019). The increase in muscle force after 4 weeks of strength training is mediated by adaptations in MU recruitment and rate coding. The Journal of physiology, 5S7(7), 1873–1887.

Del Vecchio, A., A. Holobar, D. Falla, F. Felici, R. M. Enoka, and D. Farina. “Tutorial: Analysis of MU discharge characteristics from high-density surface EMG signals.” Journal of Electromyography and Kinesiology 53 (2020): 102426.

Ditunno, J. F., Stover, S. L., Freed, M. M., & Ahn, J. H. (1992). Motor recovery of the upper extremities in traumatic quadriplegia: a multicenter study. Archives of physical medicine and rehabilitation, 73(5), 431–436.

Ditunno, J. F., Little, J. W., Tessler, A., & Burns, A. S. (2004). Spinal shock revisited: a four-phase model. Spinal cord, 42(7), 383–395.

Duan, Z., Kizyte, A., Forslund, E. B., Gutierrez-Farewik, E. M., Herman, P., & Wang, R. (2025). In vivo estimation of motor unit intrinsic properties in individuals with spinal cord injury. Journal of NeuroEngineering and Rehabilitation, 22(1), 128.

Gomes, M. M., Jenz, S. T., Beauchamp, J. A., Negro, F., Heckman, C. J., & Pearcey, G. E. (2024). Voluntary co-contraction of ankle muscles alters MU discharge characteristics and reduces estimates of persistent inward currents. *The Journal of physiology*, 602(17), 4237–4250.

Gorassini, M. A., Bennett, D. J., & Yang, J. F. (1998). Self-sustained firing of human MUs. Neuroscience letters, 247(1), 13–16.

Gorassini, Monica, Jaynie F. Yang, Merek Siu, and David J. Bennett. “Intrinsic activation of human motoneurons: possible contribution to MU excitation.” Journal of neurophysiology 87, no. 4 (2002): 1850–1858.

Gorassini, M. A., Knash, M. E., Harvey, P. J., Bennett, D. J., & Yang, J. F. (2004). Role of motoneurons in the generation of muscle spasms after spinal cord injury. Brain, 127(10), 2247–2258.

Goreau, V., Morvan, Q., Hug, F., Le Sant, G., Gross, R., & Cattagni, T. (2025). Modulation of persistent inward currents in alpha motoneurons with joint angle depends on muscle length. Journal of Neurophysiology, 134(2), 493–503.

Hassan, A. S., Fajardo, M. E., Cummings, M., McPherson, L. M., Negro, F., Dewald, J. P., … C Pearcey, G. E. (2021). Estimates of persistent inward currents are reduced in upper limb MUs of older adults. The Journal of Physiology, 5SS(21), 4865–4882.

Heckman, C. J., & Enoka, R. M. (2012). MU. Comprehensive physiology, 2(4), 2629–2682.

Hiersemenzel, L. P., Curt, A., & Dietz, V. (2000). From spinal shock to spasticity: neuronal adaptations to a spinal cord injury. Neurology, 54(8), 1574–1582.

Holstege, J. C., & Kuypers, H. G. J. M. (1987). Brainstem projections to spinal motoneurons: an update. Neuroscience, 23(3), 809–821.

Hounsgaard, J. O. R. N., Hultborn, H. A. N. S., Jespersen, B., & Kiehn, O. (1988). Bistability of alpha-motoneurones in the decerebrate cat and in the acute spinal cat after intravenous 5-hydroxytryptophan. The Journal of physiology, 405(1), 345–367.

Hultborn, H., Denton, M. E., Wienecke, J., & Nielsen, J. (2003). Variable amplification of synaptic input to cat spinal motoneurones by dendritic persistent inward current. The Journal of physiology, 552(3), 945–952.

Inglis, J. G., & Gabriel, D. A. (2021). Sex differences in the modulation of the MU discharge rate leads to reduced force steadiness. Applied Physiology, Nutrition, and Metabolism, 4C(9), 1065–1072.

Javeed, S., Zhang, J. K., Greenberg, J. K., Botterbush, K., Benedict, B., Plog, B., … C Nerve Transfers in Spinal Cord Injury (NT-SCI) Consortium. (2024). Impact of upper limb motor recovery on functional independence after traumatic low cervical spinal cord injury. Journal of neurotrauma, 41(9-10), 1211–1222.

Jenz, S. T., Beauchamp, J. A., Gomes, M. M., Negro, F., Heckman, C. J., & Pearcey, G. E. (2023). Estimates of persistent inward currents in lower limb motoneurons are larger in females than in males. Journal of neurophysiology, 129(6), 1322–1333.

Johnson MD, Kajtaz E, Cain CM, Heckman CJ (2013) Motoneuron intrinsic properties, but not their receptive fields, recover in chronic spinal injury. J Neurosci 33:18806–13.

Johnson, M. D., Thompson, C. K., Tysseling, V. M., Powers, R. K., & Heckman, C. J. (2017). The potential for understanding the synaptic organization of human motor commands via the firing patterns of motoneurons. Journal of neurophysiology, 118(1), 520–531.

Kizyte, A., Zhang, H., Forslund, E. B., Gutierrez-Farewik, E. M., & Wang, R. (2025). Neuromuscular adaptations in soleus and tibialis anterior muscles in persons with spinal cord injury. Journal of NeuroEngineering and Rehabilitation, 22(1), 239.

Koller, M. (2016). robustlmm: an R package for robust estimation of linear mixed-effects models. Journal of statistical software, 75, 1–24.

Lee RH, Heckman CJ. Bistability in spinal motoneurons in vivo: systematic variations in persistent inward currents. J Neurophysiol. 1998 Aug;80(2):583–93. doi: 10.1152/jn.1998.80.2.583. PMID: 9705452.

Lee, R. H., & Heckman, C. J. (2000). Adjustable amplification of synaptic input in the dendrites of spinal motoneurons in vivo. Journal of Neuroscience, 20(17), 6734–6740.

Lenth, R. (2023). emmeans: Estimated Marginal Means, aka Least-Squares Means_. R package version 1.8. 5.

Lüdecke, D., Ben-Shachar, M. S., Patil, I., Waggoner, P., & Makowski, D. (2021). performance: An R package for assessment, comparison and testing of statistical models. *Journal of open source software*, 6(60).

Mateo, S., Roby-Brami, A., Reilly, K. T., Rossetti, Y., Collet, C., & Rode, G. (2015). Upper limb kinematics after cervical spinal cord injury: a review. Journal of neuroengineering and rehabilitation, 12(1), 9.

Martinez-Valdes, E., Negro, F., Laine, C. M., Falla, D., Mayer, F., & Farina, D. (2017). Tracking MUs longitudinally across experimental sessions with high-density surface electromyography. The Journal of physiology, 595(5), 1479–1496.

McKay, W. B., Ovechkin, A. V., Vitaz, T. W., Terson de Paleville, D. G. L., & Harkema, S. J. (2011). Neurophysiological characterization of motor recovery in acute spinal cord injury. Spinal Cord, 49(3), 421–429.

Mesquita, R. N., Taylor, J. L., Heckman, C. J., Trajano, G. S., C Blazevich, A. J. (2024). Persistent inward currents in human motoneurons: Emerging evidence and future directions. Journal of Neurophysiology, 132(4), 1278–1301.

Murray KC, Nakae A, Stephens MJ, Rank M, D’Amico J, Harvey PJ, Li X, Harris RL, Ballou EW, Anelli R, Heckman CJ, Mashimo T, Vavrek R, Sanelli L, Gorassini MA, Bennett DJ, Fouad K (2010) Recovery of motoneuron and locomotor function after spinal cord injury depends on constitutive activity in 5-HT2C receptors. Nat Med 16:694–700.

Murray KC, Stephens MJ, Ballou EW, Heckman CJ, Bennett DJ (2011) Motoneuron excitability and muscle spasms are regulated by 5-HT2B and 5-HT2C receptor activity. J Neurophysiol 105:731–48.

National Spinal Cord Injury Statistical Center. (2024). 2024 annual statistical report for the spinal cord injury model systems (Complete public version). University of Alabama at Birmingham. https://sites.uab.edu/nscisc

Negro, Francesco, Silvia Muceli, Anna Margherita Castronovo, Ales Holobar, and Dario Farina. “Multi-channel intramuscular and surface EMG decomposition by convolutive blind source separation.” Journal of neural engineering 13, no. 2 (2016): 026027.

Oliveira, D. S. D., Carbonaro, M., Raiteri, B. J., Botter, A., Ponfick, M., & Del Vecchio, A. (2025). The discharge characteristics of motor units innervating functionally paralyzed muscles. Journal of Neurophysiology, 133(2), 343–357.

Oudega, M., & Perez, M. A. (2012). Corticospinal reorganization after spinal cord injury. The Journal of physiology, 590(16), 3647–3663.

Pearcey, G. E., & Rymer, W. Z. (2022). Population recordings of human MUs often display’onion skin’discharge patterns--implications for voluntary motor control. arXiv preprint arXiv:2208.11783.

Powers, R. K., Nardelli, P., & Cope, T. C. (2008). Estimation of the contribution of intrinsic currents to motoneuron firing based on paired motoneuron discharge records in the decerebrate cat. Journal of neurophysiology, 100(1), 292–303.

Powers, R. K., ElBasiouny, S. M., Rymer, W. Z., & Heckman, C. J. (2012). Contribution of intrinsic properties and synaptic inputs to motoneuron discharge patterns: a simulation study. Journal of neurophysiology, 107(3), 808–823.

Powers, R. K., & Heckman, C. J. (2015). Contribution of intrinsic motoneuron properties to discharge hysteresis and its estimation based on paired MU recordings: a simulation study. Journal of neurophysiology, 114(1), 184–198.

Powers, R. K., & Heckman, C. J. (2017). Synaptic control of the shape of the motoneuron pool input-output function. Journal of neurophysiology, 117(3), 1171–1184.

Sangari, S., & Perez, M. A. (2020). Distinct corticospinal and reticulospinal contributions to voluntary control of elbow flexor and extensor muscles in humans with tetraplegia. Journal of Neuroscience, 40(46), 8831–8841.

Sherrington, C. S. (1906). The integrative action of the nervous system (Vol. 35). Yale university press.

Škarabot, J., Beauchamp, J. A., & Pearcey, G. E. (2025). Human motor unit discharge patterns reveal differences in neuromodulatory and inhibitory drive to motoneurons across contraction levels. Journal of Neurophysiology, 134(5), 1429–1444.

Stephenson, J. L., & Maluf, K. S. (2011). Dependence of the paired MU analysis on MU discharge characteristics in the human tibialis anterior muscle. Journal of neuroscience methods, 1S8(1), 84–92.

Svirskis, G., & Hounsgaard, J. (1998). Transmitter regulation of plateau properties in turtle motoneurons. Journal of neurophysiology, 79(1), 45–50.

Thomas, C. K., Tucker, M. E., & Bigland-Ritchie, B. (1998). Voluntary muscle weakness and co-activation after chronic cervical spinal cord injury. Journal of neurotrauma, 15(2), 149–161.

Thomas, C. K., Bakels, R., Klein, C. S., C Zijdewind, I. (2014). Human spinal cord injury: MU properties and behaviour. Acta physiologica, 210(1), 5–19.

Thompson, C. K., & Hornby, T. G. (2013). Divergent modulation of clinical measures of volitional and reflexive motor behaviors following serotonergic medications in human incomplete spinal cord injury. Journal of Neurotrauma, 30(6), 498–502.

Tracy, B. L., Maluf, K. S., Stephenson, J. L., Hunter, S. K., & Enoka, R. M. (2005). Variability of MU discharge and force fluctuations across a range of muscle forces in older adults. Muscle & Nerve: Official Journal of the American Association of Electrodiagnostic Medicine, 32(4), 533–540.

Tysseling, V. M., Klein, D. A., Imhoff-Manuel, R., Manuel, M., Heckman, C. J., C Tresch, M. C. (2017). Constitutive activity of 5-HT2C receptors is present after incomplete spinal cord injury but is not modified after chronic SSRI or baclofen treatment. Journal of Neurophysiology, 118(5), 2944–2952.

Wiegner, A. W., Wierzbicka, M. M., Davies, L., & Young, R. R. (1993). Discharge properties of single MUs in patients with spinal cord injuries. Muscle & Nerve: Official Journal of the American Association of Electrodiagnostic Medicine, 16(6), 661–671.

Wienecke, J., Zhang, M., & Hultborn, H. (2009). A prolongation of the postspike afterhyperpolarization following spike trains can partly explain the lower firing rates at derecruitment than those at recruitment. Journal of neurophysiology, 102(6), 3698–3710.

Yamaguchi, T., Xu, J., & Sasaki, K. (2023). Age and sex differences in force steadiness and intermuscular coherence of lower leg muscles during isometric plantar flexion. Experimental Brain Research, 241(1), 277–288.

